# Predicting phage-bacteria interactions at the strain level from genomes

**DOI:** 10.1101/2023.11.22.567924

**Authors:** Baptiste Gaborieau, Hugo Vaysset, Florian Tesson, Inès Charachon, Nicolas Dib, Juliette Bernier, Tanguy Dequidt, Héloïse Georjon, Olivier Clermont, Pascal Hersen, Laurent Debarbieux, Jean-Damien Ricard, Erick Denamur, Aude Bernheim

## Abstract

Predicting how phages can selectively infect specific bacterial strains holds promise for developing novel approaches to combat bacterial infections and better understanding microbial ecology. Experimental studies on phage-bacteria interactions have been mostly focusing on a few model organisms to understand the molecular mechanisms which makes a particular bacterial strain susceptible to a given phage. However, both bacteria and phages are extremely diverse in natural contexts. How well the concepts learned from well-established experimental models generalize to a broad diversity of what is encountered in the wild is currently unknown. Recent advances in genomics allow to identify traits involved in phage-host specificity, implying that these traits could be utilized for the prediction of such interactions. Here, we show that we could predict outcomes of most phage-bacteria interactions at the strain level in *Escherichia* natural isolates based solely on genomic data. First, we established a dataset of experimental outcomes of phage-bacteria interactions of 403 natural, phylogenetically diverse, *Escherichia* strains to 96 bacteriophages matched with fully sequenced and genomically characterized strains and phages. To predict these interactions, we set out to define genomic traits with predictive power. We show that most interactions in our dataset can be explained by adsorption factors as opposed to antiphage systems which play a marginal role. We then trained predictive algorithms to pinpoint which interactions could be accurately predicted and where future research should focus on. Finally, we show the application of such predictions by establishing a pipeline to recommend tailored phage cocktails to target pathogenic strains from their genomes only and show higher efficiency of tailored cocktails on a collection of 100 pathogenic *E. coli* isolates. Altogether, this work provides quantitative insights into understanding phage–host specificity at the strain level and paves the way for the use of predictive algorithms in phage therapy.

## Introduction

Bacteriophages (phages), the viruses which infect bacteria, are ubiquitous, extremely diverse in the wild and play a key ecological role in regulating bacterial populations in various environments [1]–[3]. The interactions between phages and the bacteria they predate leads to a perpetual evolutionary arms race which is a source of major molecular innovations. As an example, recent studies uncovered more than 150 novel antiphage mechanisms used by bacteria to defend themselves against their viruses [4]. The use of phages to fight bacterial infections in a clinical context (phage therapy) has recently known a regain of interest notably due the development of antimicrobial resistance among bacterial pathogens [5], [6]. Predicting which phages precisely target and infect bacteria offers potential for the development of innovative strategies to combat bacterial infections, especially ones resistant to antibiotics [7].

Successful lytic infection of a bacterial cell by a phage is a multistep process which sequentially goes through attachment of the virion to the cell, genome injection and intracellular replication, production of new virions and release of the viral progeny through the lysis of the cell. Each step of the lytic cycle relies on specific molecular interactions involving many different bacterial and phage traits [8]. These traits encompass the bacterial hosts’ phage receptors (e.g. surface polysaccharides, membrane proteins) and its antiviral arsenal as well as their viral counterparts: phage Receptor-Binding Proteins (RBP, e.g. tail proteins) and anti-defense mechanisms (e.g. anti-CRISPR or anti-RM) [4], [9]–[11]. Those traits can be extremely diverse in natural populations, even among closely related bacteria and phages [12], [13]. Consequently, most phages are only capable of infecting hosts within a single bacterial species or genus, but even within a single species bacterial susceptibility to phages is often strain-specific [14]–[17].

Within these extremely diverse interactions, most studies on phage-bacteria interactions have focused on a few model couples (e.g. *E. coli* K-12 strain *vs.* lambda or T phages) [18]–[21]. Whether the knowledge acquired on these model organisms would be sufficient to capture a broad diversity of phage-bacteria interactions which are encountered in the wild remains to be determined.

Recent progress in bioinformatic tools to detect bacterial and phage traits from their genomes now enables a finer characterization of diverse features involved in phage-host specificity [13], [22], [23]. Predicting the phenotype of interaction between a phage and bacterial strain based solely on their respective genome thus becomes in principle more feasible [7]. Recent studies have used genomic traits to decipher phage-bacteria interactions within bacterial species including *Staphylococcus aureus* (8 phages, 259 isolates) or *Klebsiella pneumoniae* (46 phages, 138 isolates) [24], [25], paving the way to predict phage-bacteria specificity from their genomes. Current algorithms typically determine which bacterial genus a given phage can infect but cannot make predictions at deeper taxonomic levels [26]–[29]. Even within the same genus, bacterial strains may exhibit important differences in their susceptibility to phages [14], [15], [17].

*Escherichia coli* is a major pathogen worldwide, whose resistance to antibiotics remains a prime concern for public health [30]. *E. coli* belongs to the *Escherichia* genus, which is extremely diverse, notably because of its disparate ecological niches [31], [32]. The diversity of the accessory genome of *Escherichia* strains, representing more than 95% of the genus pangenome, and their phages in natural environments, highlights the wide variation existing within this genus [33].

In this work, we set out to predict phage-host specificity from genomic data in a large and diverse set of bacterial strains and phages within the *Escherichia* genus. To this end, we evaluated experimentally each individual phage-bacteria interaction between two biological collections (403 diverse *Escherichia* natural isolates and 96 phages) that we established and characterized genomically. We uncovered the traits governing phage-bacteria interactions in our dataset and designed algorithms capable of recommending phage cocktails to target specific bacterial strains within the genus *Escherichia* from a phage therapy perspective.

## Results

### Establishing a collection of diverse *Escherichia* bacterial strains and phages

To build algorithms capable of predicting the interaction between any *Escherichia* strain and any phage, we first needed an experimental dataset of interactions between a collection of *Escherichia* strains and a collection of phages. We also required that both collections – bacteria and phages – would be sufficiently large and diverse, fully sequenced and genomically characterized. To our knowledge, such a dataset was not available. We thus decided to build one *ex nihilo*.

#### The *Bertrand Picard* collection: 403 bacterial strains, representative of the *Escherichia* genus diversity

First, we put together a collection of 403 natural isolates belonging to the *Escherichia* genus that we named after the French medical bacteriologist Bertrand Picard who was a pioneer in describing the *E. coli* intraspecies diversity in relation to the lifestyle of the strains and initiated this collection in the early 1980’s (Biography in **Supplementary Text 1**). This collection is an assemblage of 369 natural isolates, which have been phenotypically well characterized in previous studies [34], [35], and of 34 strains that were used to isolate the phages in this study (**Figure 1**). A focus when establishing this collection was to encompass *Escherichia* diversity. Thus, we gathered strains from six species and eight phylogroups of the *Escherichia* genus (*Escherichia coli n* = 378). To estimate genomic diversity in a finer granularity, we also analyzed Sequence Types (ST). In the pre-genomic era, the clonal population structure of the *Escherichia* genus was typically described in terms of ST which were determined by MultiLocus Sequence Typing (MLST) [31], [36]. *In silico* MLST revealed that our collection accounts for 163 STs, with 50% of the collection covered by 14 main STs (**Figure 1A** and **1B**) [37]. Beyond ST, we analyzed additional loci known to frequently recombine such as the *rfb* locus responsible for the production of the O-antigen and the *fli* locus responsible for the production of the H-antigen. Both O- and H-antigens are surface polysaccharides which undergo strong diversifying selection and are involved in phage susceptibility [31], [32]. O and H-antigen serotypes were predicted *in silico* [37]. A total of 93 O-antigen (out of approximately 180 currently described in *E. coli*) and 41 H-antigen serotypes (out of approximately 50) are present in the *Picard* collection, including all the most abundant ones (**Figure 1C**) [38], [39]. Apart from phylogenetic diversity, we paid attention to the pathotype (commensal, environmental, pathogenic) (**Figure 1D**) and ecological source diversities of the isolates (**Figure 1E**).

**Figure 1.**
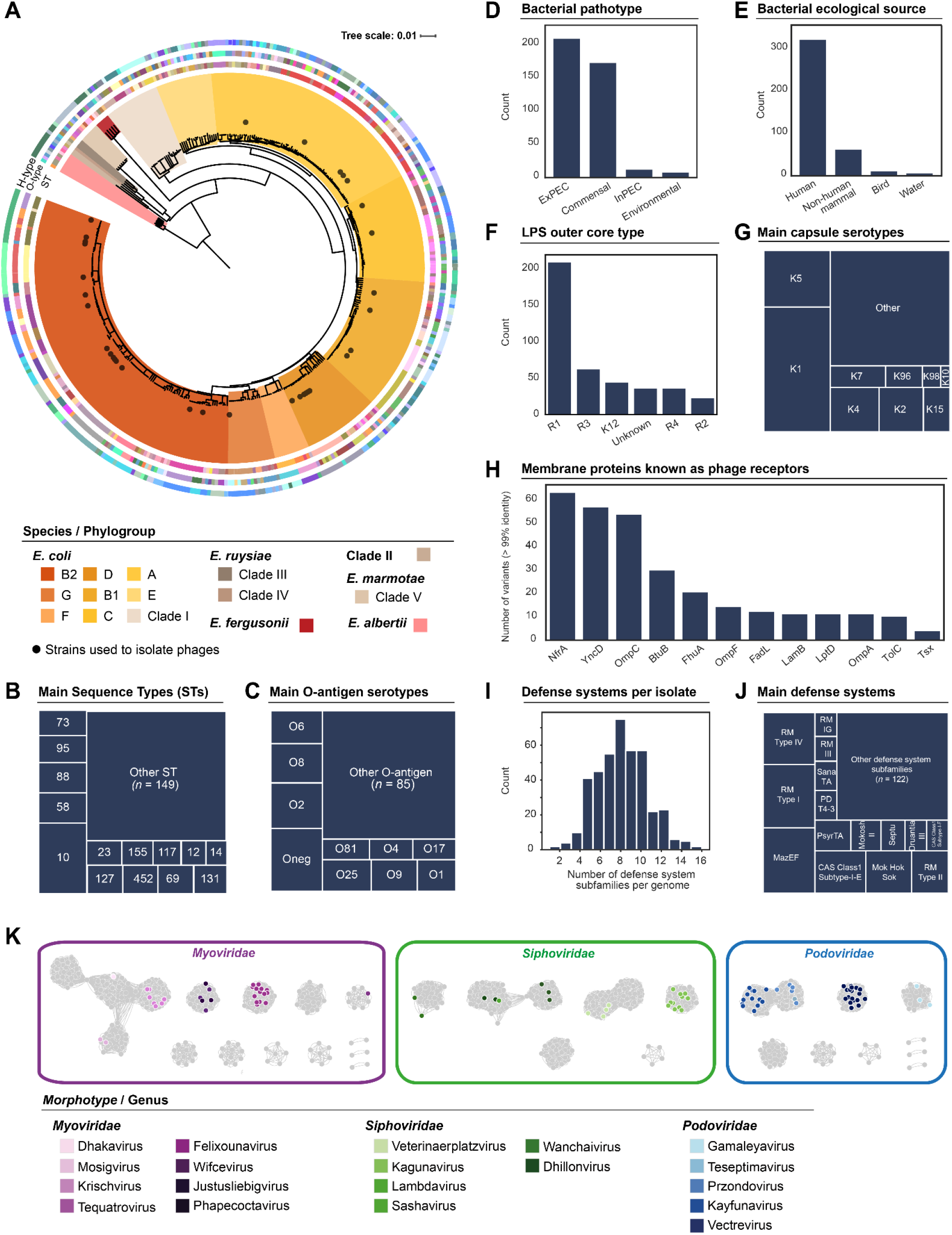
Generation of two large and diverse collections of *Escherichia* bacterial strains (*Picard* collection) and phages (*Guelin* collection). A-J The *Picard* collection of bacterial strains (*n* = 403) encompasses the phylogenetic diversity of the *Escherichia* genus and features a wide range of potential phage receptors and antiphage systems. A. Phylogenetic tree of bacterial isolates. All species of the genus *Escherichia* are present in the collection (*E. coli*, *E. fergusonii*, *E. albertii*, *E. ruysiae*, *E. marmotae* and Clade II), with representatives of all eight phylogroups (A to G) and Clade I belonging to the *E. coli* species. Each phylogroup is represented by a warm colour from brown to orange. The first ring represents the 163 different ST; the second one the 93 O-antigen types; and the third one the 41 different H-antigen types. B. Frequency of the bacterial sequence types. C. Frequency of the bacterial O-antigens. D. Number of isolates by pathotype. Collection includes clinical pathogenic isolates (*n* = 221) as well as commensal (*n* = 174) and environmental (*n* = 8) isolates. ExPEC: Extraintestinal Pathogenic *E. coli*. InPEC: Intraintestinal Pathogenic *E. coli*. E. Number of isolates by ecological source. Isolates originate mainly from human samples (feces, urine or blood, *n* = 324), non-human mammal (*n* = 61) and bird samples (*n* = 12), or water samples (*n* = 6). F. Number of isolates according to their lipopolysaccharide (LPS) outer core type. The five main LPS outer core groups are found: K-12 (*n* = 43), R1 (*n* = 206), R2 (*n* = 22), R3 (*n* = 61) and R4 (*n* = 35). Some strains could not be typed for this trait (*n* = 36). G. Frequency of the bacterial capsular serotypes (ABC capsule type). Of the 171 isolates with an ABC-dependent capsule, we identified K-antigen for 101 of them, retrieving the most described *Escherichia* capsular serotypes: K1 (*n* = 42), K5 (*n* = 19), K4 (*n* = 11), K2 (*n* = 10), K15 (*n* = 6), K7 (*n* = 6) and K10 (*n* =1). H. Number of variants of a putative phage receptor outer membrane protein. Out of the 12 outer membrane proteins previously identified as putative receptors for *Escherichia* phages [9], a large diversity of variants (gene clusters with more than 99% genomic identity) has been identified. I. Number of defense system subfamilies *per* bacterial genome. The isolates encode on average 8 defense system subfamilies, with considerable heterogeneity between strains. J. Frequency of subfamilies of bacterial defense systems. The pan immune system of *Escherichia* genus includes 137 different defense system subfamilies. K. Network representation of 1,122 *Escherichia* phage genomes. The *Guelin* phage collection (*n* = 96, colored dots) encompasses the diversity of the *Caudoviricetes* phages targeting the *Escherichia* genus. The 96 *E. coli* phage genomes in this study (in colors, current study) belong to the three morphotypes of *Caudoviricetes* (*Myo-, Podo-* and *Siphoviridae*), and cover 19 phage genera. Their taxonomic distribution was analyzed in the context of 1,026 complete genomes of reference *Escherichia* phages (in gray, *Genbank* database). vConTACT2 was used to measure the distance between phages, and Cytoscape for network visualization. Each genome is represented by a node (point); edges (line) were calculated using the number of shared protein groups. The ICTV taxonomic level for the genera of phages in the collection is represented by colored circles.

Using the genomic sequence of the strains in the *Picard* collection, we sought to characterize the traits that could potentially influence phage susceptibility. We identified the bacterial surface structures known to be potential receptors for phages [8], [9]. In *Escherichia*, many surface polysaccharides were shown to be involved in host cell recognition by phages. Out of these, the lipopolysaccharide (LPS) is a highly diverse glycolipid covering more than 75% of the bacterial outer surface. It is composed of the lipid-A, the LPS inner and outer cores, and the highly diverse O-antigen described earlier [40]. The latter two are frequently described as phage receptors in *E. coli* [9]. In our collection, we identified the five major groups of LPS outer core (**Figure 1F**). Capsules have also been previously described as receptors for capsule serotype-dependent phages in *E. coli* [41]. We developed an approach to identify strains encoding an ABC-dependent capsule and to cluster them according to their K-serotype (see methods). We observed that most of the main K-serotypes described as phage receptors were well represented in the *Picard* collection (**Figure 1G**). Capsules are well known to shape the phage host range in the *K. pneumoniae* species [25], [42] and *Klebsiella* capsules have been previously identified in *E. coli* natural isolates [43]. We hypothesized that these capsules could also influence phage susceptibility in *E. coli* and used Kaptive to detect the *Klebsiella* capsule encoding strains in our collection [44]. We found that 22 of our 403 Escherichia isolates (5.4%) encode such capsules. All the *K. pneumoniae* capsule encoding strains belonged to phylogroups A, B1 or C and had O-antigen serotypes O8, O9 or O89, in perfect agreement with a preceding study on the distribution of *Klebsiella* capsules among environmental *E. coli* isolates [43]. We also detected twelve genes coding for the outer membrane proteins that were recurrently identified in the literature as putative phage receptors for *E. coli* (**Supplementary Table 4**) [9], revealing that they are ubiquitous in *Escherichia* genomes (each one was found in more than 97% of the collection). Phage adsorption can however be easily averted by a single substitution in the gene coding for the phage receptor. When looking for the genomic diversity of genes coding for these proteins, we thus identified a large diversity of genomic variants (**Figure 1H**). Gene variants were clustered at 99% genomic identity, meaning that two variants within the same cluster diverged by at most approximately 5 amino acids from each other. Altogether these analyses provide an extensive catalog of combinations of putative phage adsorption factors within the *Picard* collection. We next focused on intra-cellular defense. Using DefenseFinder [13], we identified known antiphage systems (**Supplementary Table 5)**. On average, an isolate encodes 8 defense system subfamilies, with considerable heterogeneity among strains (i.e. from 1 to 16 defense systems) (**Figure 1I**). The pan immune system of the *Escherichia* genus comprises a total of 137 antiphage defense systems. Some of them are widespread (e.g. MazEF or RM type I), while many are present in only a few isolates (**Figure 1J**). As a result, our genomic analysis of the *Picard* collection of *Escherichia* strains provides a large panel of genomic traits relevant to investigate phage-bacteria interactions at the scale of the genus.

#### The *Antonina Guelin* collection: 96 phages, covering a large taxonomic diversity of *Escherichia* phages

We then built a collection of 96 *Escherichia*-infecting virulent phages that we named after the 20th century French physician and biologist Antonina Guelin who led the bacteriophage laboratory in the Institut Pasteur for 25 years (Biography in **Supplementary Text 2**). We isolated 94 of these phages from sewage water, collected in different locations around the Paris area (France) over a decade (see methods). The two archetypal virulent *E. coli* phages T4 and T7 were added to the 94 other phages. Importantly, the 96 phages were isolated on 34 bacterial strains which are included within the *Picard* collection (**Figure 1A**). The strains used to isolate and propagate phages cover 75% of *E. coli* phylogroups. We sequenced phage genomes and assigned functional and taxonomic annotations to each of them. All phages in the *Guelin* collection belong to one of the three *Caudoviricetes* morphotypes (*Myo-, Podo- and Siphoviridae*). This collection covers 8 subfamilies and 19 phage genera (**Figure 1K**) [45], [46].

Altogether, we built and genomically characterized two reference collections of bacterial isolates and phages specific to the *Escherichia* genus. We hypothesized that those two collections would be sufficiently large and diverse to infer generalizable rules governing phage-bacteria interactions at the strain level in the *Escherichia* genus.

### A comprehensive experimental dataset of phage-bacteria interactions for the *Escherichia* genus

Following the establishment of the *Picard* and *Guelin* collections, we assessed the susceptibility of each strain to each phage. For that, we tested each possible phage-bacteria combination. The susceptibility of the strains to phages was assessed by plaque assay experiments using three different Multiplicities Of Infection (MOI, i.e. the ratio between the number of virions and the number of bacterial cells) (MOI=10, MOI=1, MOI=0.1) (**Figure 2A**). For each phage-bacterium couple, the interaction data points were aggregated into a single interaction score. We considered that an interaction is lytic whenever we observed either individual lysis plaques or full clearing of the bacterial lawn (see methods). We chose to encode each phage-bacteria interaction into a “Minimum Lytic Concentration” (MLC) score which corresponds to the lowest concentration of phage at which a lytic interaction was observed. The MLC score is null in case of non-lytic interaction, and ranges from 1 (lytic interaction at the highest phage titer) to 4 (uncountable number of lysis plaques at the lowest tested phage titer) (see methods for details). Such encoding allows to score each interaction in a semi-quantitative way (**Figure 2B**). The final score represents the average MLC of the three independent replicates. The resulting phage-bacteria interaction matrix comprises 403 bacteria × 96 phages = 38,688 phage-bacteria interactions for the *Escherichia* genus (**Figure 2C**). We provide all associated pictures of plaque assay experiments to allow exploration of this data set in future studies (see data availability).

**Figure 2.**
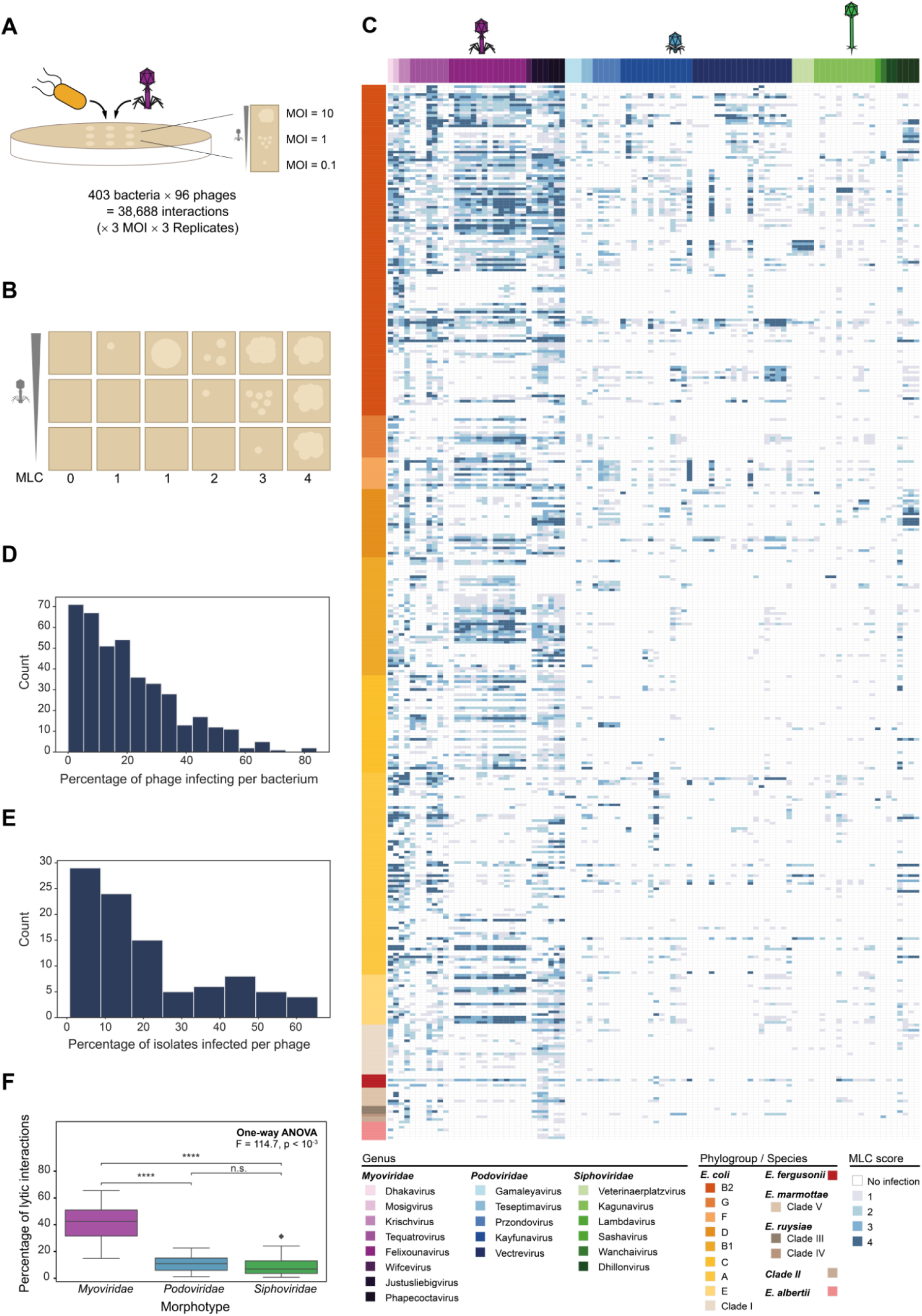
A matrix of 38,688 phage-bacteria interactions in the *Escherichia* genus reveals complex patterns. **A-B Schematics of phage-bacteria interaction outcome determination. A. Experiment set up.** We investigated the 38,688 phage-bacteria interaction combinations by plaque assay at different MOI (10, 1, and 0.1). Each experiment was performed in triplicate. MOI: multiplicity of infection. **B. Schematic of the assignment of MLC scores.** Each column corresponds to an example of how the MLC score was determined. Phage-bacteria interactions were encoded using the MLC that corresponds to the lowest concentration of the phage at which a lysis of the bacterial lawn was observed. 0 indicated that no lytic interaction was observed; 1 is lytic interaction at the highest phage titer (MOI 10); 2 is lytic interaction at middle phage concentration (MOI 1); interactions observed at the lowest MOI of 0.1 are distinguished between 3 (individualized lysis plaque) and 4 (entire lysis of the bacterial lawn) (see methods). MLC: minimum lytic concentration. **C. Phage-bacteria interaction matrix.** Each column is a phage. Each line is a bacterial isolate. Colors correspond to phage viral genera and bacterial phylogroups. Phage and bacteria are distributed according to their phylogenetic proximity as determined in figure 1. Intensity of blues in the matrix corresponds to the MLC score. **D. Percentage of phages infecting per bacterium**. The graph represents the number of bacterial isolates stratified by the percentage of the 96 bacteriophages capable of infecting them (per 5% phage susceptibility). **E. Percentage of isolates infected by each phage**. The graph represents the number of bacteriophages stratified by the percentage of the 403 isolates they can infect (per 5% phage host range). **F. Breadth of the phage host range according to the phage morphotype.** Whiskers indicate mean and 95% confidence intervals.

We first evaluated the propensity of any phage to infect any bacteria. Overall, 20.3% of the interactions were lytic. Each phage was able to infect an average of 83 (± 70) bacterial strains (20.6% of the *Picard* collection) (**Figure 2D**), while a bacterial isolate could be lysed by 20 (± 15) phages (20.8% of the *Guelin* collection) on average (**Figure 2E**). Only 12 (3.0%) isolates were not infected by any of the phages, including 5 *Escherichia* isolates not belonging to the *E. coli* species. We thus observed significant discrepancies among groups of phages and groups of bacteria. While our knowledge of phage-bacteria interactions in *Escherichia* is essentially based on model pairs of bacteria (K-12, *Escherichia* B) and phages (T phages), less than half of the phages were able to infect *Escherichia B* strains (e.g. BE: *n* = 45 (46.9%), BL21: *n* = 48 (50.0%)) and 35.4% *E. coli* K-12 (MG1655: *n* = 34 (35.4%)) (**Supplementary Figure 1**). The three phage morphotypes exhibited very different infection patterns: *Myoviridae* had a significantly broader host range than *Podo*- and *Siphoviridae* (41.3% of lytic interactions on average *vs.* 11.1% and 9.1% respectively (One-way ANOVA: F = 114.7, *p* < 10^-3^, **Figure 2F** and **Supplementary Text 3**). The full matrix exhibits an overall nested pattern (**Supplementary Text 4** and **Supplementary Figure 2**).

Overall, the global analysis of the interaction matrix shows that a broad collection of phages, characterized by their diversity, can cause a lytic interaction in 20% of phage-bacteria interactions at the strain level within the *Escherichia* genus.

### Isolation strains heavily correlate with the phage host range, likely *via* the determination of the phage RBPs

We then wanted to use this experimental dataset to predict phage-bacteria interactions. A straightforward approach would be to use the 38,688 phage-bacteria interactions which were experimentally assessed as a training dataset for a machine learning model aiming at predicting the outcome of the interaction for any phage-bacterium couple. The input traits which are *a priori* relevant for such algorithms are the adsorption factors of the phage (i.e. its receptor-binding proteins; RBP) and of the bacterial strain (i.e. all surface polysaccharides and outer membrane proteins), and the bacterial defense systems. However, the diversity of both adsorption factors (e.g. > 90 serotypes of O-antigen,) and defense system families in the collection (>100 families) makes the number of candidate traits provided as input features to the models considerable. We thus decided to reduce the number of features in order to limit the risk of overfitting our models and improve their ability to generalize to new phage-bacteria interactions. This implied to better understand what would be the traits of phages and bacteria which best explained interaction patterns in the matrix. We chose to follow a targeted approach consisting in (i) characterizing the traits seemingly relevant to study the phage-bacteria interactions at the strain level, (ii) assessing the statistical association between each genomic trait and the interaction patterns observed in the matrix, and (iii) quantifying the relative importance of each trait in explaining the infection patterns.

First, we examined phage infection patterns in the matrix. We found that phages from the same viral genus (>70% genomic identity) share more similarity in their infection patterns than what would be expected for two phages taken at random (PERMANOVA, adjusted R^2^=0.13, *p* < 10^-5^) [47]. We further observed that the major explanatory variable of phage host range is its isolation strain (PERMANOVA, adjusted R^2^=0.53, *p* < 10^-5^). The interaction between both variables accounted for more than 60% of the variance in the distance matrix (Permutational ANOVA, adjusted R^2^=0.65, *p* < 10^-5^) (**Figure 3A**) [47]. We then investigated which specific genomic traits could explain phage infection. Comparative genomics between phages of the same genus showed that most of the core genes were highly conserved in terms of amino acid identity within each genus, while genes encoding for tailspike or tail fiber proteins were systematically hotspots of variability [48]. Additionally, we observed that those tailspike and tail fiber-encoding genes were more conserved among phages isolated on the same strain, compared to phages isolated on different isolation strains (**Figure 3B** and **Supplementary Figure 3**). Tail fiber and tailspike genes are well known to encode for RBP [8], [49], [50]. We thus hypothesized that RBP could be the major phage traits explaining the infection patterns observed in the interaction matrix.

**Figure 3.**
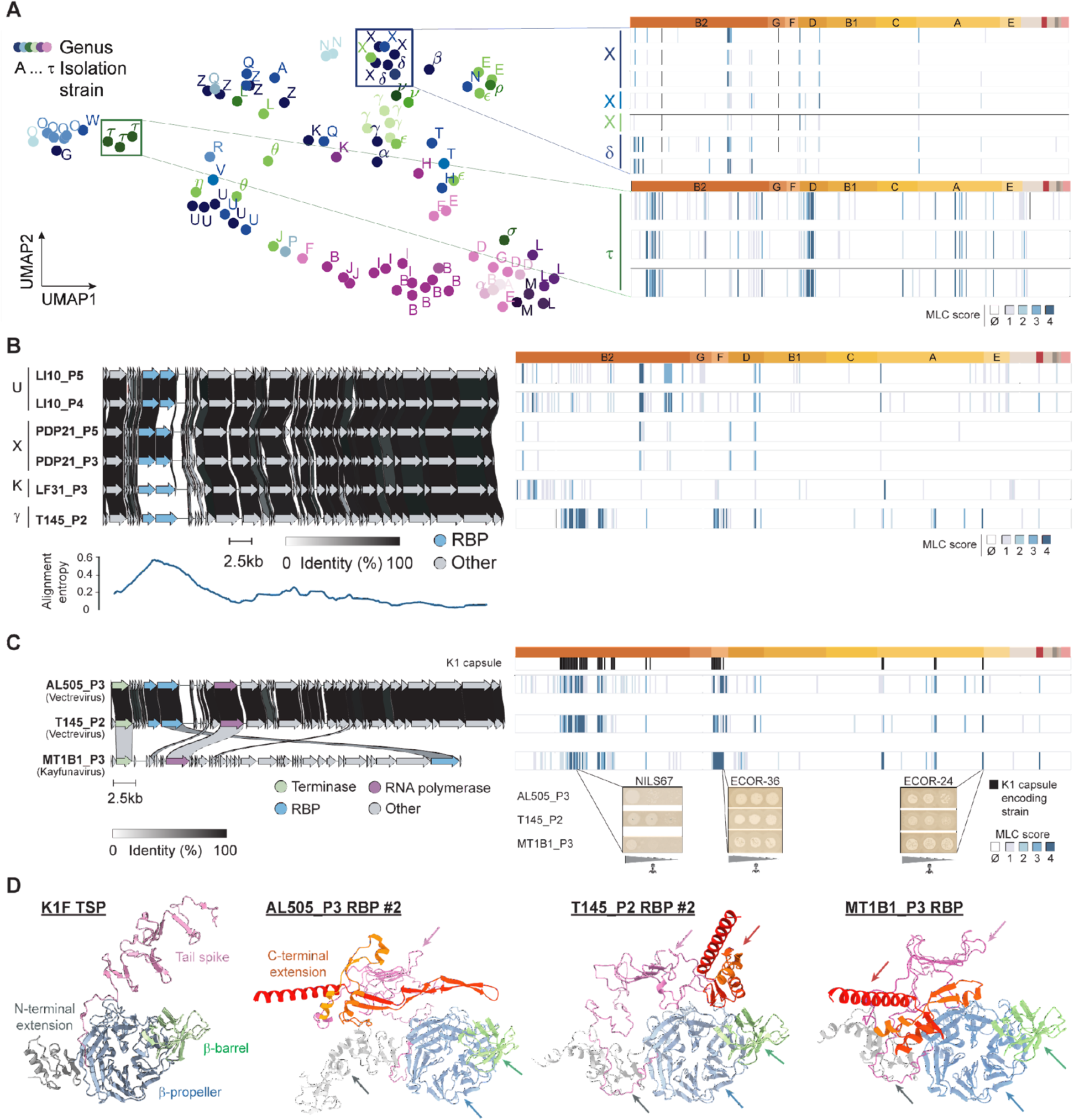
Phage isolation strain and Receptor-Binding Proteins are major determinants of the phage host range. **A. Phage genus and isolation strain associate with phage host range. Left:** Dimensionality reduction of the phage host range using the Uniform Manifold Approximation and Projection (UMAP) algorithm [55]. The 403-dimensional vector encoding the binary interactions of each phage with all the bacteria is reduced to a 2-dimensional vector. The color of each phage indicates its genus and the letter indicates the bacterial strain on which it was isolated and propagated. **Right:** Host range of the phages located within the two zoom boxes present on the UMAP graph. Phages are clustered according to their host range similarities. Shades of blue indicate the MLC score assigned to the interaction between each phage and each bacterium (Ø: No infection). **B. Phage RBP are hotspots of genomic variability and are more conserved among phages isolated on the same host. Top Left:** Genomic map of six representative phages of the *Vectrevirus* genus generated using Clinker [56]. The six phages were isolated on four different strains indicated by a letter. Genes are colored according to their functional annotation: either RBP (blue) or other (gray). The stroke between each pair of genes at the same locus indicates the percentage of aminoacid sequence similarity. **Bottom Left:** Position-wise entropy of the genomic alignment. A multiple whole genome alignment is performed including the six phage genomes and the Shannon entropy is computed at each position of the alignment based on the distribution of the nucleotides. The lower the entropy, the stronger is the conservation of nucleotides at a given position in the multiple sequence alignment. **Right**: Host range of the six phages presented in the genomic map. **C. RBP sharing is sufficient to explain host range similarity among phages even among different genera. Left:** Genomic map of the three selected phages exhibiting similar host ranges [56]. AL505_P3 and T145_P2 belong to the same genus (*Vectrevirus*) whereas MT1B1_P3 is a *Kayfunavirus*. Genes which are conserved (> 30% amino acid similarity) are colored according to their functional annotation. **Right:** Host range of the three phages. Strains which encode in their genome an ABC-dependent capsule with serotype K1 are indicated (black). For the three phages, plaque assay raw images are provided on three example bacteria (NILS67, ECOR-36 and ECOR-24) at the three MOI. **D. T145_P2, AL505_P3 and MT1B1_P3 share a beta-propeller containing RBP also found in the K1F phage.** Predicted structure of the monomer of the conserved RBP among AL505_P3, T145_P2 and MT1B1_P3 [52]. The RBP contains an endosialidase catalytic beta-propeller domain (PF12217, light blue) containing a nested beta-barrel (residues 104-543, light green) upstream of a beta-prism/tailspike domain (residues 544-718, pink) and variable N- and C-terminal extensions (gray and red). The X-ray diffraction structure of the K1F phage tailspike (TSP) monomer is also shown as a reference (PDB: 3GVJ). The name of each structural domain is given on the example of K1F TSP and its position in AL505_P3 RBP #2, T145_P2 RBP #2 and in MT1B1_P3 RBP is indicated by arrows.

To explore this hypothesis, we identified a triplet of phages which exhibit very similar infection patterns while belonging to different genera and being isolated on three distinct isolation strains (**Figure 3C**). We sought out to identify the genetic factors underlying the similarity in their infection patterns and wondered if this similarity could be explained by shared RBP. Two of these phages (T145_P2 and AL505_P3) belong to the *Vectrevirus* genus (*Autographiviridae/Molineuxvirinae*) but the third one (MT1B1_P3) is a *Kayfunavirus* (*Autographiviridae/Studiervirinae*) and shared very few genomic similarity with the other ones. The three phages shared only three of their core genes: (i) their large terminase subunit, (ii) their RNA polymerase which are also conserved among all *Vectreviruses* and *Kayfunaviruses* and (iii) a gene identified as a RBP in the *Kayfunavirus* MT1B1_P3 and in both *Vectreviruses*. While sharing limited amino acid identity (56% between the *Vectrevirus* and *Kayfunavirus* cognate gene), domain annotation using the PFAM-A database and the structure prediction of each RBP monomer allowed us to identify that the three RBP of interest share a common architecture (**Figure 3D**) [51], [52]. Structural homology search revealed that this protein architecture can be found in other phage tailspikes such as in the K1F phage tailspike (K1F TSP) which have been shown to use K1-serotype capsule as a target [49], [53], [54] (**Figure 3D**). K1F TSP has been shown to bind to polysialic acid and have and endosialidase activity [54]. We hypothesized that such RBP could therefore also use K1 capsules as a receptor. We identified 45 strains encoding a K1-serotype capsule in the *Picard* collection including 4 isolation strains, 3 of which were precisely the isolation strains of phages T145_P2, AL505_P3 and MT1B1_P3. When testing the decision rule that our three phages of interest infect exactly the K1-encapsulated strains, we obtained a precision of 73% and a recall of 72%. As a comparison, we applied the same decision rule – lytic interaction of a bacterial strain if and only if the strain harbors a K1-serotype capsule – to all the other *Vectreviruses* and *Kayfunaviruses* and obtained a precision of 7% and a recall of 14%. This suggests that T145_P2, MT1B1_P3 and AL505_P3 exhibit important similarities in their infection patterns because they possess a similar RBP in spite of being isolated and propagated on distinct bacterial strains. This detailed example further shows that phages which share their RBP, regardless of their isolation strain, also have similar infection patterns. Altogether, this further strengthens the idea that RBP are the major determinant of phage host range. The systematic identification of RBP in all our phages also allowed us to show that the number of lytic interactions performed by a phage was not linked with the number of RBPs it encodes, but rather likely to be influenced by the targeting specificity of its RBPs (**Supplementary Text 5**).

Overall, our general statistical analysis supports the idea that phages isolated on the same host exhibit similar infection patterns because they tend to share similar RBP. As such, the diversity of isolation strains plays a fundamental role when selecting phages with a diverse set of RBPs. This implies that the choice of the isolation strains is an important parameter to take into account when isolating new phages, especially in the context of phage therapy.

### Bacterial adsorption factors drive their susceptibility to phages

We then investigated the association of bacterial traits with infection patterns in the matrix and paid particular interest to core phylogeny, adsorption factors and antiphage defense systems. First, we observed that bacterial strains which are phylogenetically closely related exhibit some limited but significant correlation in their host range (Mantel test, *r=*0.03, *p*=2×10^-3^) [47]. However, this significance was lost whenever pairs of strains with a phylogenetic distance below 10^-4^ substitutions per position were removed (typically less than a few hundred SNPs on the whole core genome), indicating that the correlation was driven by very tightly related kins (**Supplementary Figure 4**). Our dataset shows that phylogeny poorly explains phage susceptibility.

We next investigated which bacterial traits exhibit the strongest association with the bacterial infection patterns, notably trying to evaluate the relative importance of adsorption and intracellular defense. To do so, we assessed the association of bacterial adsorption factors and defense systems with the probability of being infected by each individual phage. This analysis was performed within the framework of Linear Mixed-Effects models to handle the phylogenetic non-independence between bacteria (see methods). We fitted one Generalized Linear Mixed-Model (GLMM) per phage, aiming at explaining the lytic interactions of its corresponding phage based on both the adsorption factors and the defense systems subfamilies [17], [57]. A total of 30 bacterial adsorption factors significantly associated with phage lytic interactions whereas only 2 defense systems did the same (**Figure 4A**). We also followed a complementary statistical approach which gave similar results (**Supplementary Text 6**). This suggests that, similarly to what is observed in phages, bacterial adsorption factors were the best variable to explain the overall infection patterns.

**Figure 4.**
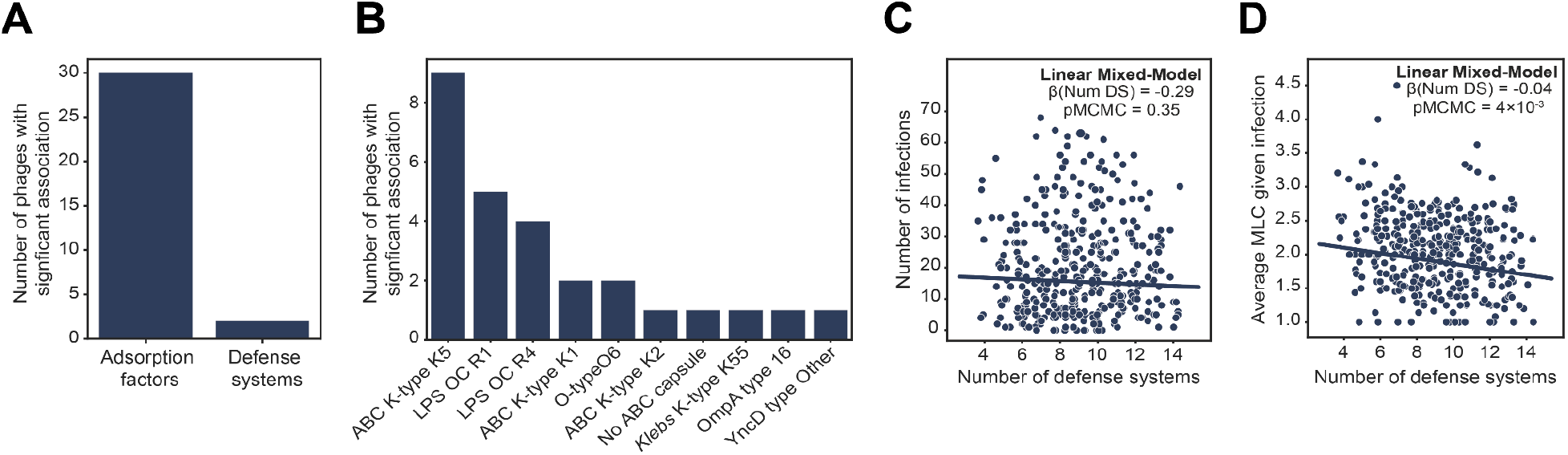
Bacterial adsorption factors are the major determinants of phage-bacteria interactions in contrast with defense systems which marginally reduce bacterial susceptibility to infecting phages. **A. Number of bacterial adsorption factors and defense systems significantly associated with phage lytic interaction.** A distinct binomial GLMM is fitted on the lytic interactions performed by each phage. GLMM are provided with bacterial adsorption factors (O-antigen serotypes, H-antigen serotypes, ABC-dependent capsular serotype, *Klebsiella* capsular serotype, outer membrane protein types) and defense systems subfamilies as covariates. We then count how many times each category of bacterial traits (adsorption factor or defense system) significantly associates with the lytic interaction by any phage. We consider that the association is significant if pMCMC is lower than 10^-3^ (see methods) [57]. **B. Number of times each individual bacterial adsorption factor associates significantly with phage lytic interaction.** As in **B**, one binomial GLMM is fit on the interactions of each individual phage with the bacterial phylogeny as a random effect [57]. Here, each model takes only the bacterial adsorption factors as covariates. We then count how many times each bacterial trait associates significantly (pMCMC < 10^-3^) with phage lytic interaction. K-type: ABC dependent K-antigen serotype. LPS OC: LPS Outer Core. O-type: O-antigen serotype. **C. Number of infections undergone by each bacterial isolate as a function of the number of antiviral defense systems detected in the genome of the strain.** The best regression line obtained by fitting a LMM using the number of defense systems as a covariate and taking the core bacterial phylogeny into account as a random effect is shown [57]. The mean of the posterior distribution obtained for the regression coefficient associated with the number of defense systems β(Num DS) as well as the estimated pMCMC are reported. **D. Average MLC observed for each strain when taking only the lytic interactions undergone by the strain into account as a function of the number of antiviral defense systems detected in its genome.** As in **C.**, the best fit line obtained by fitting a Linear Mixed-Model taking the number of defense systems as a covariate and the core bacterial phylogeny as a random effect is represented. The mean of the posterior distribution as well as the probability of the coefficient being null under the posterior are also reported.

We then sought out to characterize more precisely which adsorption factors would associate the most with phage infection in the matrix. To do so, we fitted a distinct GLMM on each individual phage infection patterns, taking only the bacterial adsorption factors as covariates. Here, we identified a total of 27 adsorption factors significantly associated with the lytic interactions of 24 phages in the *Guelin* collection. The most frequent adsorption factors yielded by the linear models were mostly surface polysaccharide-related traits (K-serotype, LPS outer core or O-antigen serotype). Interestingly, outer membrane proteins – frequently identified as a phage receptor in several studies – were not associated with phage infection in our dataset [9], [58] (**Figure 4B**). Importantly, specific adsorption factors such as the *Klebsiella* capsule serotype exhibited a significant but weak quantitative association with the infection patterns in the matrix (PERMANOVA, adjusted R^2^=0.02, *p<*10^-3^). We argue that in spite of their weak quantitative association with infection patterns, those traits are worth investigating as they highlight local, non trivial, patterns in the matrix. Phage PDP110_P1, for example, infected only three bacteria in the full *Picard* collection including its isolation strain (0.7% of lytic interactions for this phage) (**Supplementary** Figure 5A). Strikingly, the three infected bacteria were scattered across phylogroups A and C and matched exactly the three strains encoding a *Klebsiella* capsule with serotype K55. Furthermore, raw plaque assay results showed that each infection event led to the apparition of halos surrounding lysis plaques which is commonly mentioned as an indicator of surface polysaccharide degradation by enzymes expressed by the phage (**Supplementary Figure 5B**) [25]. This strongly suggests that phage PDP110_P1 is a K55-targeting phage.

### Defense systems are not required to predict phage-bacteria interactions in our dataset

As we were surprised by the limited explanatory power of antiphage systems on the host range in our statistical inference models, we further investigated their impact on phage-bacteria interactions. First, we evaluated if bacteria encoding more defense systems would be protected against more phages. We found no evidence of such association (LMM, β_*Num DS*_ = −0.22, 95% credible interval = [-0.74, 0.34], pMCMC=0.41) (**Figure 4C**) [57]. We further hypothesized that bacterial antiphage systems function once a phage has adsorbed and is starting the replication cycle. As such, defense systems could interfere with the ability of an infecting phage to multiply in the cell, instead of impacting the set of phages which can infect the cell. This would imply that an increase in the number of defense systems encoded by a strain would not impact the number of infecting phages, but rather would make infecting phages less virulent (i.e. with a decreased MLC score). To test this hypothesis, we assessed the correlation between the number of defense systems encoded in a strain and its average MLC upon phage infection (taking only the lytic interactions into account in the regression). We observed a significant but low correlation between larger defense arsenal and decreased MLC given phage infection (LMM, _β*Num DS*_ = −0.04, 95% credible interval = [−0.05, −0.01], pMCMC=4×10^-3^) (**Figure 4D**). This correlation was robust to the removal of extreme values and to the random subsetting of bacterial ST indicating that it was not driven by a limited number of bacterial clades. To control for potential confounding factors that would be associated with antiphage systems, we also assessed the statistical association between the average MLC given infection and (i) the number of antimicrobial resistance (AMR) genes encoded in each bacterial genome or (ii) the bacterial genome size. We chose these controls because they follow similar evolutionary dynamics as the number of defense systems and we *a priori* do not expect them to correlate with the average MLC given infection. Both controls strengthened the idea that the negative correlation between the number of defense systems encoded by a strain and its average MLC given infection is not spurious (see **Supplementary Text 7** and **Supplementary Figure 6**). It is important to note that although significant, the effect size of the number of defense systems on the MLC given infection was low (β_*Num DS*_ = −0.04). Those results strongly suggest that, in our collection, the number of defense systems of a bacterial strain does not influence the probability of infection by phages in our interaction dataset but could slightly impact the level of susceptibility to already infecting phages. Overall, these results show that bacterial defense systems are not required to predict the phage-bacteria interactions and can be removed from the set of candidate traits provided as input features to our models.

### Adsorption factors are sufficient to accurately predict phage-bacteria interactions at the strain level

Identifying the bacterial and phage genomic traits allowed to dramatically narrow down the number of traits that we should provide as input features to a model aiming at predicting phage-bacteria interactions at the strain level for phage-bacteria couples in the *Escherichia* genus. Given the results of the statistical inference (see previous paragraph), we hypothesized that the adsorption factors alone would be sufficient to predict the phage-bacteria interactions. To test this hypothesis, we trained machine learning models to predict the phage-bacteria interactions as a binary classification task (i.e. lytic *vs.* non lytic interaction). A distinct model was used for each individual phage based only on the bacterial adsorption factors and the bacterial core phylogeny (no data regarding defense systems was provided as inputs to the models). The performance of each phage-specific model was evaluated using Group 10-Fold Cross-Validation. In this setup, bacterial strains are clustered according to their phylogenetic relationships such that two closely related strains will always be grouped together, either in the train or in the validation set. Here, we chose to group bacteria into the same block if their core phylogenetic distance was lower than 10^-4^ substitutions per position. This allows to evaluate the capacity of the model to generalize beyond a certain phylogenetic distance, potentially to completely new ST. Importantly, all the interactions of a given bacterial strain are also always grouped together in the train or in the validation set, forcing the model to learn to predict phage-bacteria interactions based solely on genomic traits (in this specific case, the adsorption factors and the core phylogeny) and not based on previously observed interactions. Phage-specific models achieved an average Area Under the ROC curve (AUROC) of 77% (**Figure 5A**). When aggregating each phage-specific model prediction, an overall AUROC of 86% was obtained on Group 10-Fold Cross-Validation (**Figure 5B**). This was particularly true for phages with as few as 10% of lytic interactions which indicates good generalization ability even for very specific phages (**Figure 5C**). This holds true for weakly infecting phages exhibiting strong class imbalance (e.g. 33% of our phages infect less than 10% of the bacteria in the *Picard* collection), a phenomenon known to hinder the training of traditional machine learning algorithms [59]. Our detailed understanding of which phage-bacteria interactions are correctly predicted provide a roadmap for further research as it will allow us to focus on the 3,379 false negatives and 2,922 false positives that are not captured by our current genomics-based predictive model over a total of 38,688 interactions (**Figure 5D**). The prediction performance we achieve shows that the adsorption factors are sufficient to accurately predict phage-bacteria interactions in our dataset.

**Figure 5.**
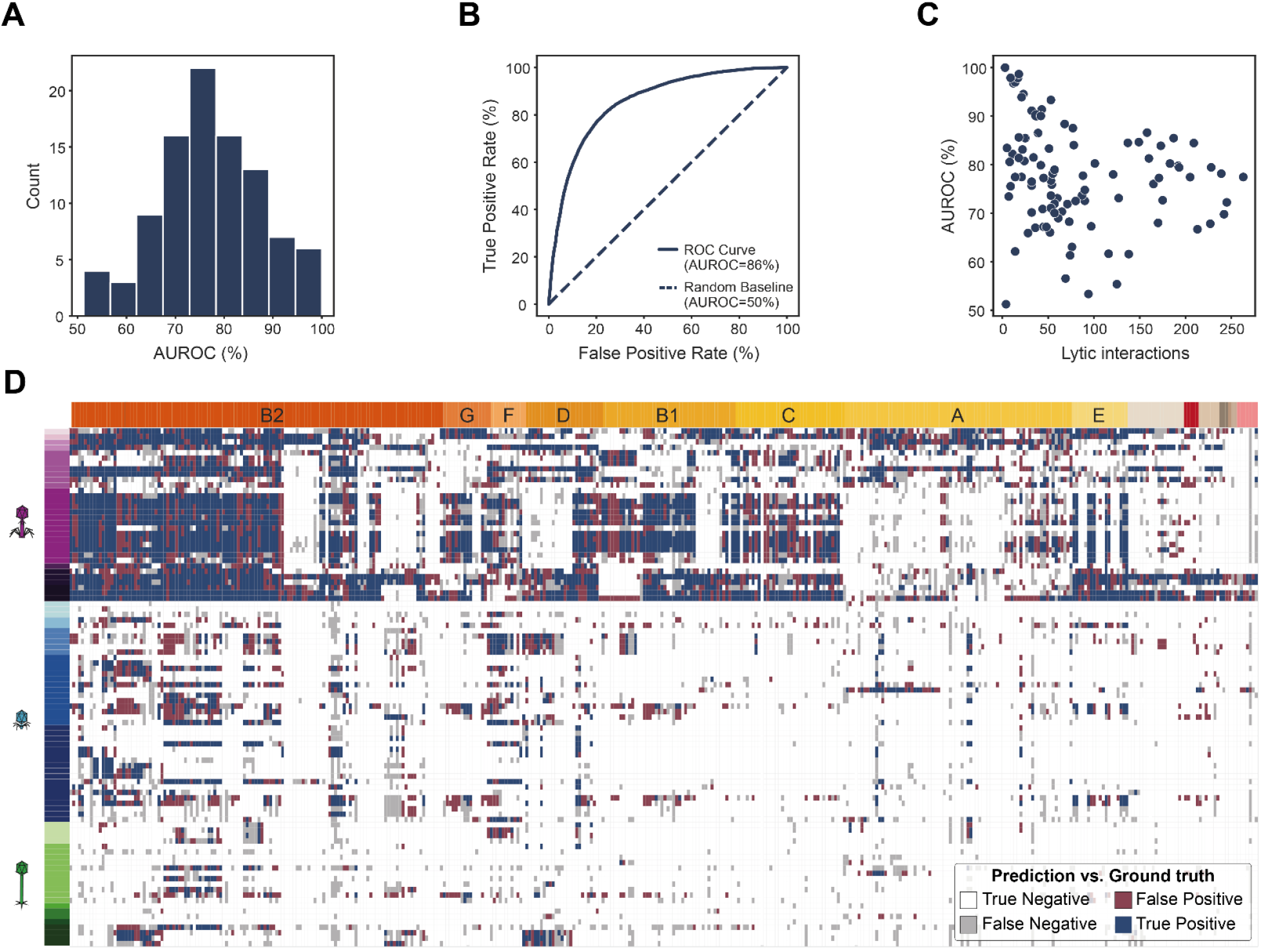
Adsorption factors are sufficient to accurately predict phage-bacteria interactions, even for phages with very few lytic interactions. **A. Distribution of the prediction performance among phage-specific models.** One classification model is fitted for each individual phage and evaluated on Group 10-Fold Cross-Validation. The performance on the validation sets of each phage-specific model is measured by the AUROC metric and averaged across all cross-validation folds. By definition, AUROC ranges from 50% (random predictor) to 100% (perfect predictor). **B. Overall prediction performance of the predictive algorithm across the 96 phages.** Contrary to what is presented in **A.** the predictions of each phage-specific model on the validation set at each cross-validation fold are aggregated together. Prediction performance of the obtained model is assessed using the ROC curve and the AUROC score (blue). **C. Distribution of the prediction performance among phage-specific models as a function of the phage number of lytic interactions.** As in **A.**, the performance of each phage-specific model on the validation sets of each cross-validation fold is averaged. It is plotted against the number of lytic interactions performed by its corresponding phage, which gives the amount of class imbalance in the classification problem. **D. Prediction matrix obtained using adsorption factors and core phylogeny only.** One classification model per bacteriophage is trained to predict the phage-bacteria interactions bacteriophage from the interaction matrix. Each model takes as input the same bacterial adsorption factors as well as the core phylogeny of the bacteria. They are trained and evaluated on Group 10-Fold Cross-Validation. Predictions obtained on the validation set at each round of the cross-validation procedure are compared with the ground truth and are reported in this prediction matrix. Blue: True Positive. Red: False Positive. Gray: False Negative. White: True Positive.

### Genomic traits allow to successfully recommend tailored phage cocktails to target pathogenic bacterial strains

We showed that the detected genomic traits hold predictive power regarding phage host range and that it was possible to train machine learning models predicting interactions for any phage-bacteria couple with satisfying performance. We next evaluated whether our predictive models for phage-bacteria interactions could be used in a framework of phage therapy where a combination of several virulent phages is assembled into a phage cocktail to target a specific pathogenic bacterial strain (hereafter referred to as the “query” strain) [60]. We aimed at designing a machine learning-based phage cocktail recommender system to target pathogenic *E. coli* strains based on bacterial genomic traits only and to evaluate the performance of the algorithm *in vitro* on a test collection of previously unseen *E. coli* pathogenic isolates (**Figure 6A**).

**Figure 6.**
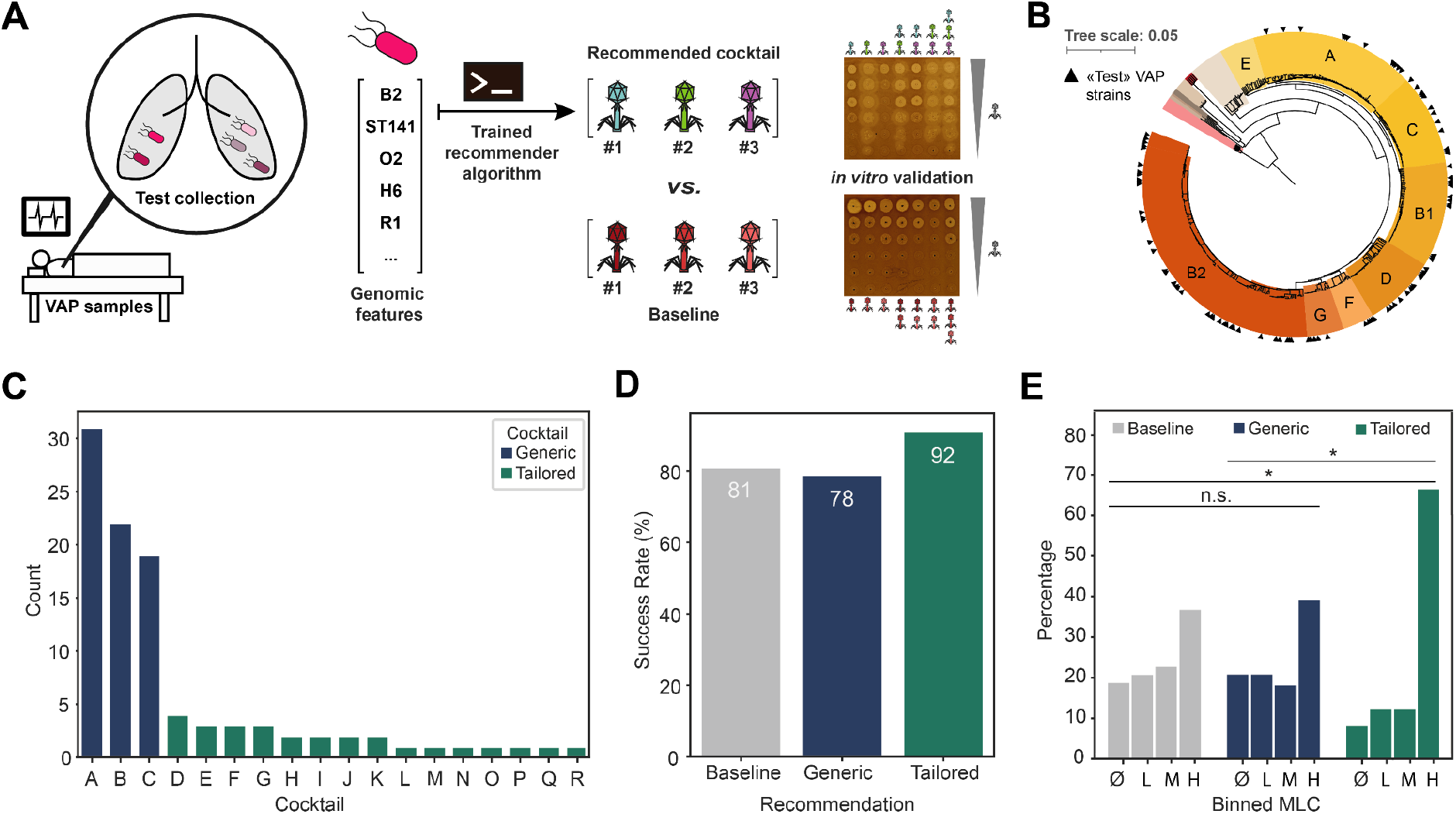
A recommender system allows the design of phage cocktails targeting previously unseen pathogenic bacteria based on their genomic traits. **A. Schema of the experiment.** A new «test» collection of 100 *E. coli* pathogenic strains is isolated from clinical samples of Ventilator-Associated Pneumonia (VAP). Each test strain is sequenced and genomically characterized to identify the genomic features of interest. A recommender algorithm is trained on the original interaction matrix to predict the interaction between a query bacterium (pink) with any phage of the *Guelin* collection based solely on its genomic traits. Phages are then sorted according to their predicted probability of infection and a cocktail of three phages is recommended. The recommended cocktails are then evaluated by challenging *in vitro* each strain in the test collection with its recommended cocktail as well as with a baseline cocktail composed of the three most covering triplet of phages in the original interaction matrix from six different MOI (from 10 to 0.0001). **B. Phylogenetic tree of the 100 strains of the test collection together with the 403 strains of the *Picard* collection.** The test collection is constituted of 100 *E. coli* strains sampled from the *ColoColi* collection of isolates. Core phylogeny of the 100 test strains is computed jointly with the 403 strains of the *Picard* collection. Phylogroups of the *E. coli* strains are specified by their name. Strains of the test collection are identified by a black triangle. **C. Distribution of the 18 different cocktails recommended to the 100 *E. coli* strains in the test collection.** Each cocktail is named with a letter from A to R. Recommended cocktails are categorized into either «generic» or «tailored» according to the steps of the pipeline at which they were recommended. **D. *In vitro* success rate of the recommended cocktails on the test collection**. Success rate is the percentage of lytic interactions performed by each category of cocktail (Baseline, Generic, Tailored) over the total number of times such a cocktail category was recommended. **E. Binned MLC obtained *in vitro* on the test collection.** The MLC score (ranging from 0 to 6) is binned into four ordered categories: Ø («No infection», MLC=0), L («Low», 1 ≤ MLC ≤ 2), M («Medium», 3 ≤ MLC ≤ 4) and H («High», 5 ≤ MLC ≤ 6). The distribution of the binned MLC for each cocktail type (Baseline, Generic, Tailored) is then measured on the test collection. Asterisks indicate a statistically significant difference among cocktail groups in terms of MLC (two-sided Mann-Whitney test, p < 0.05), n.s.: not significant. MLC: minimum lytic concentration.

We first set out to determine what should be the size, *i.e.* the number of phages, in any cocktail. This parameter should be large enough to cover diverse bacteria and provide more robustness to erroneous phage recommendations but remain low to avoid undesired side effects and limit the cost of production. We evaluated the number of infected bacteria by the most covering *k*-uplet of phages. We observed that the most covering triplet of phages infects 63% of the *Picard* collection whereas the most covering 4-uplet of phages only reached 68% of infected bacteria (**Supplementary** Figure 7). This small increase in efficiency suggested that a cocktail size of *k=3* phages offered a good trade-off between efficiency and parsimony, leading us to choose this value for the cocktail size in the rest of the experiments.

We then went on to design a recommender system that goes sequentially through four steps and stops whenever three phages are recommended. At each step, the probability of infection for each phage of the *Guelin* collection is computed. Phages are ranked according to their predicted probability of infection and top-ranked phages are added to the recommended cocktail when their probability is higher than 0.5. The four steps detailed in the methods section are 1) Whether the query strain is similar to an isolation strain (similarity is computed using haplotype (ST/O-antigen serotype/H-antigen serotype) and adsorption factors), 2) whether the query strain has the same haplotype to other strains from the *Picard* collection, 3) whether the query strain has similar adsorption factors to other strains from the *Picard* collection, 4) whenever three phages could not be recommended in the upstream steps, the available places in the cocktail filled with the most generalist phages in the *Guelin* collection (**Supplementary** Figure 8). The rationale behind this pipeline organization of the recommender algorithm was to prioritize steps which were empirically measured as more precise but often less exhaustive over steps which are more exhaustive but less precise.

We then assessed the capacity of our recommender system to generalize to a new collection of 100 previously unseen *E. coli* isolates. Those 100 “test” strains were selected from the *ColoColi* collection which contains 310 *E. coli* natural isolates collected from bronchoalveolar samples of patients and responsible for Ventilator-Associated Pneumonia (VAP) (**Figure 6A**) [61]. The test strains were chosen to be as diverse as possible in terms of phylogeny (60 different ST, 26 of which were represented in the *Picard* collection) and geographical origin (**Figure 6B** and **Supplementary Table 6**). STs which were typical of VAP such as ST127 or ST141 were kept in the test strains. Each of these natural isolates was sequenced and genomically characterized using the same procedure as described before. Phage cocktails were recommended based solely on the genomic traits of these 100 strains. A total of 18 different phage cocktails were recommended for the whole test collection. We observed that the recommended cocktails could be gathered into two categories based on the steps of the pipelines at which phages were recommended. First, some cocktails (*n*=15, recommended to 24 test bacteria) contain phages recommended at early stages of the pipeline (denoted as “tailored” in the rest of the text) while the rest of the cocktails (*n*=3, recommended to 76 bacteria) are recommended at late stages of the pipeline (denoted as “generic” in the rest of the text) (**Figure 6C)**. This behavior is a direct consequence of the pipeline organization of our recommender algorithm and reflects the fact that some bacteria – those which recommended tailored cocktails – are easily handled by the pipeline whereas others are handled with more difficulty and tend to be recommended generic cocktails. As a consequence, tailored cocktails sometimes contained highly specific phages (e.g. T145_P2 which infects 13.1% of the original collection) often recommended early in the pipeline compared with generic cocktails which only contained broad range phages (56% of lytic interactions on average on the *Picard* collection) often recommended downstream in the pipeline (**Supplementary** Figure 9). To evaluate the performance of our algorithm, we compared the performance of the tailored and generic cocktails with an uninformed baseline cocktail. The latter corresponds to the most covering triplet of phages, i.e. the cocktail of the three phages which together lyse the largest number of bacteria in the original interaction matrix. This baseline cocktail yielded productive lysis on 63% of the original *Picard* collection.

We then evaluated experimentally the tailored, generic and baseline cocktails by plaque assay experiments. To do so, we tested each possible decomposition of the cocktails (each phage individually, each pair of phages and the full 3-phages cocktail) in order to identify putative synergistic or antagonistic behavior between phages. Interestingly, no antagonistic nor synergistic phage-phage interaction was observed in this experimental setup. To evaluate the performance of the cocktails, we first compared the overall percentage of lysed bacteria for each method. The baseline and the generic recommendations had similar percentage of success (81.00% and 78.95% lysed bacteria) whereas the tailored cocktails had a higher success rate on the bacteria it was recommended to (91.67% lysed bacteria) (**Figure 6D**). Second, we evaluated the MLC of the full cocktail. The tailored cocktails significantly outperformed both the baseline (two-sided Mann-Whitney test: U=399.5, *p*=0.02) and the generic cocktails (two-sided Mann-Whitney test: U=1171.5, *p*=0.02, **Figure 6E**). Overall, our algorithm successfully recommended tailored phage cocktails based only on genomic data. The recommended cocktails could complement a broad baseline approach in a typical clinical use.

## Discussion

Our work demonstrates that it is feasible to predict phage-host specificity from genomes to a certain extent. With over 38,688 interactions from 403 *E. coli* isolates and 96 phages, we understand 65% of the variance in phage host range based on the identified phage genomic traits. We predict phage-bacteria lytic interactions from bacterial genomic traits with an AUROC of 85%. We further show that such predictions can be harnessed to design tailored efficient phage cocktails.

Our predictions could be improved and refined. Our collections, while large, only represent a subset of the diversity of all *Escherichia* natural isolates. Some adsorption factors were not fully identified in our research, including the Enterobacterial Common Antigen (ECA) or the Wzy-dependent capsular serotypes in *E. coli* [9], [62]. Additional less targeted statistical approaches such as GWAS could be used to uncover associations by infection patterns and other bacterial traits. Genomic analyses also have inherent limitations, as they do not consider gene regulation and expression in response to the bacterial environment. The expression of many proteins including certain outer membrane proteins or capsular components may exhibit variability depending on the prevailing bacterial environment [63], [64]. We evaluated phage-bacteria interactions under controlled conditions, specifically in a non-limited medium. Different environmental conditions could induce distinct surface phenotypes in bacteria, potentially resulting in entirely disparate outcomes in their interactions with a particular phage.

Despite these limitations, our results provide a clear picture on the genomic determinants of phage-host specificity in our collection of natural *E. coli* isolates: adsorption factors play a major role and antiphage systems a marginal one. These results question the ecological role of defense systems. Previous work on *Vibrio crassostreae* showed that very closely related strains with different defense systems are susceptible to different phages [14], [15]. However, analyses on our interaction dataset cannot identify any significant correlation between the number of antiphage systems and the number of phages infecting a given isolate. This might be a consequence of the evolutionary scale at which our collection is established. When looking at the whole genus level as we do here, bacterial adsorption factors are shuffled among strains and might appear as the primary determinant of bacterial susceptibility to phage. When studying at a finer evolutionary scale (e.g. among strains sampled from the same location at short time intervals), the effect of defense systems might become quantitatively more important as their turnover rate is faster than the one of adsorption factors [65]. In our collection, we were able to associate a higher number of bacterial defense systems with a reduced virulence of infection. However, the observed effect size is low. This observation could be explained by a low probability that a random defense system acts against a specific phage, backing the potential existence of a “pan-immune system” (natural bacterial communities are heterogeneous and individual strains encode only a small subset of all the defense potential of the community). Alternatively, this could also suggest that these antiphage systems hold an alternative role, such as competitions between very diverse MGEs as suggested by recent literature [66], [67]. Whether this holds true for other bacterial genera, remains to be determined.

Beyond providing quantitative answers to the contribution of different drivers of phage-bacteria interactions, our results generate a roadmap to move forward in the understanding of these complex interactions. First, our work demonstrates the diversity of phage-bacteria interactions beyond laboratory models (e.g. MG1655 *vs.* T phages). By making available to the community two new collections of diverse phages and bacterial isolates, and the accompanying interaction dataset for each phage-bacteria interaction pair available through the Viral Host Range database [68], we hope to provide a starting point for exploring the biology of such interactions in a more mechanistic manner. Second, as we showed in **Figure 4E**, genomic-based predictions are also useful to point out the specific phage-bacteria interactions we poorly understand and might require further attention. Both the depth and breadth of the matrix and the genomic and phenotypic characterization allow to map the existing knowledge efficiently and can thus orient future studies to the interactions not explained by our current models. We also believe that employing the same systematic approach could greatly enhance our comprehension of phage-bacteria interactions in different bacterial species like other ESKAPE pathogens. This approach may unveil previously undiscovered intricacies specific to each bacterial species.

Finally, we used predictions to inform phage therapy applications. Our understanding of a vast dataset of phage-bacteria interactions demonstrates that the strain used to isolate phages has a dramatic impact on phage tropism and is a powerful experimental leverage to increase the diversity of phage host range of a phage collection. We took advantage of our dataset of interactions to train a phage cocktail recommender system. Our results indicate that it is possible to recommend cocktails to previously unseen *E. coli* pathogenic strains with satisfying performance. When recommending phage cocktails using our pipeline, the clinician could provide the genome of a pathogenic bacterial strain in the pipeline and either get a prediction for a “tailored” cocktail, which is likely to have a higher success rate and a stronger lysis of the query bacterium or if no tailored cocktail can be predicted a generic one should be used. An algorithm designed to aid decision making based on genomic data could streamline and expedite the selection of phages and tailor cocktails for enhancing the efficiency of phage therapy while minimizing potential side effects, in a standpoint of personalized medicine.

## Methods

### Culture media and solutions

Strains were cultivated at 37°C on LB Lennox medium (agar or broth) (Becton Dickinson, USA). The thickness of the agar plates was standardized using an automatic pump dispensing 50 mL per 120mm x 120mm petri dish on a flat surface. Phosphate-buffered saline (PBS) 1x was prepared by dissolving commercial tablet (Sigma-Aldrich, USA) in 200 mL of deionized water and sterilized by autoclaving. All the batch media were prepared within a few days to ensure the most homogeneous condition.

### *Picard* collection (bacterial strains)

The *Picard* collection includes 369 strains, previously gathered to be representative to the phylogenetic diversity of the *Escherichia* genus and described by Galardini et al. [69], and the 34 strains used to isolate and propagate the phages were also added to this collection. All the details of the 403 strains are available in the **Supplementary Table 1.**

### *ColoColi* collection (bacterial strains)

To test the validity of our predictive phage cocktail algorithm, in a context compatible with phage therapy settings, we used a collection of 100 pathogenic strains of *E. coli*. These strains originated from the *ColoColi* collection, which have been previously described [61]. In brief, *ColoColi* gathered 310 strains mainly from pneumonia acquired under mechanical ventilation (VAP) from 14 different hospital centers in France. It includes in majority pathogenic strains (*n* = 263) and some colonizing strains (*n* = 47). Half of the strains used to isolate and propagate phages of the *Guelin* collection came from the *ColoColi* collection. Sampling of this collection led to the selection of 100 strains, excluding the phage host strains, and representing 7 phylogroups of *E. coli*. No phylogroup H strains or cryptic clades were present in the *ColoColi* collection. These strains represented 60 different STs, had 32 different O antigens and 24 H antigens. All the details of the 100 strains are available in the **Supplementary Table 6.**

### *Guelin* collection (bacteriophages)

Phages were isolated from samples of sewage water collected over 10 years from several wastewater treatment plants of the urban region of Paris (France). Phage isolation was performed on a specific host with an enrichment step as described by Debarbieux et al. [70]. Isolation host strains were all *E. coli* isolates, including clinical isolates from ventilator associated pneumonia (*n* = 21), urinary tract infection (*n* = 2), and intestinal strains from inflammatory bowel disease patients (*n* = 7), as well as EAEC (*n* = 1), commensal strain of mice (*n* = 1) and lab strains (K-12 and *E. coli* B, *n* =2). All phage lysates were filtered (0.2 µm) before use. By convention, phage titer was determined on the host strain used to isolate and propagate the phage. Phages closely related to other ones were deliberately excluded if they shared a nucleotide identity upon 98% unless they displayed obvious phenotypic differences of their host-range. Ultimately, out of 128 isolated phages, among which few were previously characterized [71]–[77], 96 unique phages were obtained. All the details of the 96 bacteriophages are available in the **Supplementary Table 2.**

### Assembling phages in cocktails

Three phages which have been predicted by the recommender algorithm to be able to lyse the query strain were assembled in a cocktail, and compared to a reference “Baseline” cocktail, composed of the three phages whose combined cover the wider host range of the *Picard* collection (LF82_P8, CLB_P2, LF73_P1). For both baseline cocktail and predicted cocktails, the phages have been tested individually (phage 1, phage 2, phage 3), in two-by-two combinations (phages 1 & 2, phages 1 & 3, phages 2 & 3) or all-together (phages 1, 2 & 3) following the same plaque assay experimental method described below. Each combination of two or three phages was a mix of an equivalent quantity (Plaques Forming Unit, PFU) of each phage, and so as to obtain phage solutions with final theoretical concentrations of 5.10^8^ PFU/mL. The quantity for each phage was calculated from the concentration of the phage determined on its own isolated strain. The composition of phage cocktails was detailed in the **Supplementary Table 7**.

### Evaluating phage-bacteria interaction outcomes: plaque assays experiments

The infectivity of a given phage on a given bacterial strain was quantified by plaques assays. The phage-bacteria interaction was assessed by adding a 2 µL drop of phage solution on a bacterial lawn on an agar plate. Each bacterial strain was overlaid on agar plates, from a fresh liquid medium culture, that reached the exponential growth phase (OD_600nm_ 0.25 ≈ 5.10^7^ CFU/mL). All the phage solutions titers were normalized at 5.10^8^ PFU/mL in PBS. Each phage against each bacteria was tested at different MOI (number of phage/number of bacteria). The multiplexing and standardization of phage deposit was performed by automation with a Viaflo 96 device (Integra Bioscience, Swiss). Agar plates were incubated overnight at 37°C. The reading of the results was standardized. All plates were digitized with a high-resolution scanner after 16 hours of incubation and reviewed manually using a custom interface built on top of the Napari interactive viewer (v0.4.16) [78]. The result of each interaction was encoded according to defection or not in the bacterial lawn. We encoded each phage-bacteria interaction using the “Minimum Lytic Concentration” (MLC) score, which corresponds to the lowest concentration of the phage at which a lytic interaction is observed. We consider that an interaction is lytic whenever we observe individual lysis plaques or full clearing at any MOI. Individual lysis plaques ascertain the lysis of the bacterial population with production of new virions. Clearing of the bacterial lawn at high MOI could result from productive lysis of the bacterial population by the phage, or from another mechanism such as lysis from without, abortive infection. The average MLC upon infection was computed by taking only the lytic interactions of a given bacterial strain and averaging the MLC score across all infecting phages.

Three independent replicates were performed to assay the interactions of the 38,688 phage-bacteria couples (403 isolates from the *Picard* collection versus the 96 bacteriophages from the *Guelin* collection), at three different MOI: 10, 1, 0.1 or 0.001, corresponding to phage concentrations of 5.10^8^ PFU/mL (replicates R1, R2 and R3), 5.10^7^ PFU/mL (replicates R2 and R3), 5.10^6^ PFU/mL (replicates R1, R2 and R3) and 5.10^4^ PFU/mL (replicate R1). The MLC score is null in case of non-lytic interaction, and ranges from 1 (lytic interaction at the highest phage titer) to 4 (uncountable number of lysis plaques at MOI 0.1). The outcome of interaction at MOI 0.001 was not taken into account in the calculation of the MLC score because it was not verified by a replicate. We measured that 98.35% of the interactions – regarding any phage/bacteria/MOI triplet – yielded consistent annotated scores across triplicates (either same score assigned to the three replicates (88.64%) or same score assigned to 2/3 replicates (9.71%)).

Three independent replicates were performed to assay the interactions of the phages of the cocktails (alone, or in combination) with the 100 isolates from the *ColoColi* collection, at six different MOI: 10, 1, 0.1, 0.01, 0.001, 0.0001 corresponding to phage concentrations of 5.10^8^ PFU/mL, 5.10^7^ PFU/mL, 5.10^6^ PFU/mL, 5.10^5^ PFU/mL, 5.10^4^ PFU/mL and 5.10^3^ PFU/mL. The MLC score is null in case of non-lytic interaction, and ranges from 1 (lytic interaction at the highest phage titer) to 6 (uncountable number of lysis plaques at the lowest phage titer).

### Genome sequencing

Phage nucleic acid was extracted by a conventional molecular method. Briefly, after a pre-treatment of the lysate with DNase and RNase, capsid was disrupted by adding proteinase K and SDS, and the nucleic acid retrieved by phenol-chloroform-acid-isoamyl alcohol 25:24:1 separation followed by a sodium acetate-ethanol precipitation step. Bacterial nucleic acid was extracted using the EZ1 DNA tissue kit (Qiagen, Germany) on EZ1 Advanced XL (Qiagen, Germany) according to the manufacturer’s recommendations. The libraries were constructed using the Nextera DNA prep Kit (Illumina, USA) according to the manufacturer’s protocol. A pair-end sequencing (300 bp) was performed on a MiniSeq platform (Illumina, USA).

### Genome analysis

Read quality was checked using FastQC [79]. Adapters and quality reads were trimmed by TrimGalore. Reads were *de novo* assembled with SPAdes v3.15.4 [80] in one single contig. Ambiguities were resolved for phage genomes with the Bandage visualizing tool. We used Prokka v1.14.5 (--kingdom Viruses) [81] to annotate the genomes. All the genome sequences have been deposited on the ENA under the Bioproject ID: PRJEB69283.

### Phylogenetic analysis

*In silico* typing of the bacterial isolates was carried out using SRST2 for the sequence type (ST) according to the MultiLocus Sequence Typing (MLST) Warwick. Phylogroup was confirmed by the Clermont Typing software [82]. Bacterial genomes were annotated and compared with PanACoTA v.1.3.1 [83] and phylogenetic trees were annotated with iTOL v5 [84] online tool.

Phage were sorted by family and genus based on whole genome BLASTn using Viridic v1r3.6 [85] compared against the latest release of the ICTV classification VMR_MSL38_v2 (September 2023). Virus assignment into genera (≥70% similarities) and species (≥95% similarities) ranks follows the International Committee on Taxonomy of Viruses (ICTV) genome identity thresholds.

For the network analysis, distance between phage were calculated by vConTACT2 v0.11.3, using both viral genomes of this study as well as the 1,026 *Escherichia* phage complete genomes from *Genbank* (Organism name containing “Escherichia” OR “Coli’’ in all viruses genomes taxid 10239). Phage taxonomic network was then visualized and annotated with Cytoscape software v3.9.1 [86].

### Detection of genomic traits from the genomes of phages and bacteria

Antiphage defense systems were detected using DefenseFinder v1.2.0 using DefenseFinder models v1.2.3 with default parameters [13], CapsuleFinder v.1 with default model for the ABC-dependent capsule except for the parameter inter_gene_max_space which was set to 20 [87]. We also detected and typed the ABC-dependent capsules to known ones using custom MacSyFinder models (available in the github repository of the project) [22]. Those models were created by clustering (at 80% identity and 80% coverage) all the genes present between the boundaries of ABC capsules detected by CapsuleFinder in 1843 *Escherichia coli* complete genomes from RefSeq using mmseqs2 v13.45111 [88]. Colocalization of the different clusters was then used to form capsule type definitions and the sequences were used to create custom HMM profiles. HMM profiles and MacsyFinder models are available (see data availability). The attribution of known capsules type was done using those models on reference K-antigen serotype loci. The *Klebsiella* capsular serotypes were determined using Kaptive v.2.0.6 with default parameters and the Kaptive reference database for *Klebsiella* K loci [44].

### Methods regarding Nestedness and Modularity of the interaction matrix

The nestedness temperature (T) and the modularity score (Q) of the interaction matrix were assessed using respectively the BINMATNEST algorithm [89] and the BRIM algorithm [90]. Null distributions for T and Q were obtained by generating *n*=1,000 random matrices with identical shape and average connectance (probability that a given point in the matrix yields a lytic interaction) as the original interaction matrix. T and Q scores were assessed for each random matrix and the p-value for T and Q in the original matrix was computed from the null distribution.

### Statistical inference regarding phage genomic traits

To investigate differences in host range breadth among the three phage morphotypes present in the *Guelin* collection, we performed an ANOVA followed by Tukey honest significance test.

To explore the links between phage traits and the phage host range in a multivariate approach (i.e. taking all phage-bacteria interactions into account at once), we built a distance matrix for each phage-phage pair based on their respective binary infection vector (403 components for each phage). Since most of the phage-bacteria interactions are non-lytic, the Jaccard distance was used as a metric in order to only take lytic interactions into account when comparing two phages. The obtained pairwise distance matrix was then used to (i) perform exploratory data analysis using dimensionality reduction with the UMAP algorithm [55] and (ii) partition the distance matrix between diverse sources of variations using PERMANOVA [47]. In brief, PERMANOVA is a nonparametric alternative to ANOVA that allows to partition variance in a distance matrix between diverse groups defined by a covariate of interest (e.g. the phage genus), and to test if the fitted coefficients are significantly different from zero using permutations of the elements in the distance matrix. We performed PERMANOVA as implemented in the *adonis2* function of the vegan R package (v2.6-4) [47].

To identify which genes are responsible for encoding RBPs, we first generated a set of candidate genes using the algorithm published by Boeckaerts et al. [91]. We then manually curated this list of candidate genes using several approaches. For phage subfamilies which contained a well-studied phage (e.g. *Tevenvirinae* and T4), we exploited the conservation of gene synteny. For the rest of the phages, we performed (i) domain annotation using HMMER (version 3.3.2) and the PFAM-A database (version 34.0) and (ii) structure prediction using ESMFold to identify if a given protein was likely to function as a RBP. HMMER hmmsearch function was run with the --cut_ga option. Functional annotation and structure prediction also allowed us to determine whether a given RBP was a tailspike (typically containing a beta-helix domain) or a tail fiber (long, unstructured monomer) [8], [49], [51], [52]. The accurate identification of the RBP encode in each phage genome allowed us to investigate host range similarity between three phages, T145_P2, AL505_P3 and MT1B1_P3. RBP structural similarity analysis was performed using the Foldseek Search server [53].

### Statistical inference regarding bacterial genomic traits

The correlation between phylogenetic distance and host range similarity in bacteria was assessed with the Mantel test as implemented in the vegan R package (version 2.6-4) [47]. In this analysis, only *E. coli* isolates were taken into account to properly assess the within-species association between phylogeny and host range. Pairwise distance matrices were computed for the bacterial core phylogeny using PanACoTA v1.3.1. Gene families were defined at 80% identity when computing the pangenome and a gene family frequency of 100% when computing the core genome [83]. Pairwise distance was also computed for bacterial binary infection vectors (96 components in each vector) using the Jaccard distance (see previous paragraph for additional information) and the Mantel test was applied to the two matrices using 10,000 free permutations and the Spearman rank correlation coefficient.

The relative importance between bacterial adsorption factors (surface polysaccharides, outer membrane proteins) and intracellular defense systems to explain phage infection was assessed using a univariate approach (*i.e.* by analyzing the infection patterns of one specific phage at a time). We chose to use Linear Mixed-Effects models to perform this analysis. LMM allows to perform inference in cases where observations are not independent, e.g. when they come from a split-plot design or when they share an evolutionary history as is the case in this study. To not take this phylogenetic non-independence between bacteria could lead to inflated type I error rate (more false positive detections of significant associations). As the response variable is binary (non lytic *vs.* lytic interaction), we used a Generalized Linear-Mixed Model (GLMM) which extends the framework of the linear mixed-effects model to non normally distributed response variables. We fitted one binomial GLMM per phage using the MCMCglmm R package v2.35 [57]. To do so, we included both the bacterial adsorption factors and the defense systems as covariates. Only traits with more than 8 observations were included in the models in order to avoid model overparameterization. The inverse relatedness matrix computed from the core phylogenetic distance matrix was also included in each model as a random effect to account for phylogenetic non-independence among bacteria. Each phage specific model was fitted for a total of 500,000 iterations, a burning time of 50,000 iterations and a thinning value of 250 iterations. Again, each model was fitted using an inverse Gamma prior (V=1 and ν=2×10^-3^) and convergence was assessed by checking the effective sample size and parameter autocorrelation. The significance of a statistical association was assessed using the provided pMCMC associated with each trait. pMCMC is the probability of 0 under the posterior distribution estimated for each coefficient. We chose a significance threshold of 10^-3^.

We also followed a complementary approach which relied on multivariate statistical models. The rationale behind this approach was to treat infection patterns in the matrix as whole, instead of separating phages during the analysis as was in the univariate approach. We fitted 3 PERMANOVA models: the first one took both adsorption factors and defense systems as covariates (“full” model), the second took only adsorption factors as covariates and the third one took only antiphage defense system subfamilies as covariates. All three models using the vegan R package (v2.6-4) were fitted for 10,000 iterations and adjusted R^2^ was estimated using the sequential sum of squares. Phylogenetic non-independence of bacteria was taken into account by including in the models eight covariates which were computed from the core phylogenetic tree of bacteria using Principal Coordinates Analysis (PCoA). Those eight covariates were introduced as the first covariates of the models so that the phylogenetic signal they encode was taken into account before the effect of additional covariates (e.g. adsorption factors or defense systems) was estimated. Finally, the three models were compared based on the Akaike Information Criterion (AIC) which values both goodness of fit and parsimony (number of degrees of freedom) of a statistical model. The lower the AIC, the better is the model fit given a number of parameters.

To further investigate which adsorption factors could be explain the infection by each individual phage, we fitted another GLMM per each phage taking all the bacterial adsorption factors as covariates, the bacterial core phylogeny as a random effect and trying to explain its corresponding phage infection patterns. Each model was fitted using an inverse Gamma prior (V=1 and ν=2×10^-3^), 500,000 iterations, a burning time of 50,000 iterations and a thinning parameter of 250 iterations. Again, traits with less than 8 observations were lumped to avoid model overparameterization and a significance threshold of 10^-3^ was chosen.

Finally, we estimated the statistical association between either the number of defense systems encoded in each bacterial, the genome size of the strain or the number of AMR genes it encodes, and the number of lytic interactions it undergoes or its average MLC upon infection. Average MLC upon infection is computed by taking all the lytic interactions observed for a strain and averaging the MLC of each infecting phage. This metric measures the “strength” of an infection given that a lytic interaction occurs, regardless of whether the bacterial strain is infected by many phages or not. Each statistical association was estimated using a simple LMM since the target variable (either the number of lytic or the average MLC upon infection) is real valued and is normally distributed. Each model was provided with the inverse relatedness matrix computed from the bacterial phylogeny as described in previous paragraphs and was fitted using for 1,000,000 iterations, with a burning time of 50,000 and a thinning parameter of 250 iterations. Model convergence was checked as described earlier. Estimated posterior mean for the intercept and slope were used to draw the best fit line shown in **Figures 4C**, **4D** and in **Supplementary** Figure 6.

### Training algorithms to predict phage-bacteria interactions

After having identified genomic traits which overall associate with the infection patterns in the matrix, we sought to train predictive algorithms for the phage-bacteria interactions. We chose to predict a binary response variable i.e. whether a given phage-bacterium couple will give a lytic interaction or not as this problem formulation makes it simpler to design and evaluate predictive algorithms. For each phage, four classification models were trained. The four models were (i) Logistic Regression with L2 penalty, (ii) Logistic Regression with L1 penalty, (iii) Random Forest Classifier with a maximum depth of 3 and 250 decision stumps (iv) Random Forest with a maximum depth of 6 and 250 decision stumps [92]. To treat the problem of class imbalance, a class weight was parameterized according to the percentage of positive (lytic) interactions performed by this phage. The weight of the negative class was always set to 1. For phages with more than 60% of positive interactions, the weight of the positive class was set to 0.8; for phages with between 40% and 60% of positive interactions it was set to 1; between 30% and 40% of positive interactions positive class weight was set to 1.5; between 20% and 30% of positive interactions, class weight was set to 2 and below 20% of positive interactions, class weight was set to 3. All models were provided with the same input features: bacterial core phylogeny and bacterial adsorption factors. Bacterial core phylogeny was embedded using the UMAP algorithm into 8 coordinates [55] and adsorption factors were one hot encoded. To ease model training, adsorption factor sharing of each bacterium with the host strain of the phage was also explicitly encoded into features. To avoid model overparameterization, categorical features with less than 3 observations were lumped. All models were fitted with the scikit-learn python module version 1.1.2. Each fitted model was evaluated using Group 10-Fold Cross-Validation in which train and validation sets were split following bacterial blocks defined by core genome similarity. In our setup, bacteria with a core phylogenetic distance of less than 10^-4^ substitution per position were grouped together in the same block. For each phage, the average AUROC obtained on the validation sets of each Cross-Validation were compared and the best model was kept.

### Building a recommender system for phage cocktails

The recommender system was designed to contain four successive prediction steps. At each step, all phages were ranked based on their predicted probability of infecting the query regarding a specific criterion/decision rule and the top-ranked phage(s) were recommended. The first step (“Similar as isolation strain”) aimed at comparing the query strain with the isolations strains in the *Picard* collection based on their adsorption factors and haplotype (ST/O-antigen serotype/H-antigen serotype). The specific decision rules to compare the query strain with isolation strains were automatically learned by a single random forest algorithm (max depth of 4; 80 estimators and a class weight of 1.5 for the positive class). Phages were ranked according to their predicted probability based on this criterion, and any phage with a predicted probability higher than 0.5 was added to the cocktail. The second step (“Similar as strains in the *Picard* collection”) of the pipeline compares the query strain to the rest of the strains in the *Picard* collection based on their haplotypes, using a *k-*neighbors-like approach: if a group of strains with the same haplotype as the query was identified, phages which infect the majority of the bacteria in this group were recommended. The third step compared the query with the strains of the *Picard* collection using the adsorption factors and a random forest classifier as in the first step (max depth of 4; 80 estimators and a class weight of 1.5 for the positive class). The fourth step was reached if the algorithm did not manage to recommend three phages upstream in the pipeline. A naive strategy was then adopted: to fill the remaining places in the cocktail with the most generalist phages (highest number of lytic interactions) in the *Guelin* collection. The rationale behind this step is that if the algorithm failed to recommend algorithms regarding the criteria implemented upstream in the pipeline, it indicated that the learned decision rules were not exhaustive enough to recommend phages to this specific query strain. During the process, we explicitly forbid two phages from the same genus and isolated on the same host to be recommended to a given query strain. This design rule guarantees some diversity in the phage cocktail and, in the case of erroneous predictions, avoids recommending three similar which will fail together. When the implemented decision rules do not capture the query strain, the only rational decision is to rely on the best uninformed decision rule we have, i.e. recommending phages which lyse the highest number of bacteria in the training set similar to what was done to choose the baseline cocktail.

## Acknowledgements

We are grateful to members of the MDM lab, Beatriz Beamud, David Bikard, Ariel Lindner and Vincent Libis for their useful comments on earlier versions of the manuscript. We thank Damien Maura, Mathieu Galtier, Marine Henry, Nicolas Dufour, Raphaëlle Delattre and Marta Lourenço from the Bactériophage, Bacterium, Host lab of Institut Pasteur for the isolation of phages of the *Guelin* collection. We thank the ColoColi group for sharing the strains of the *ColocColi* collection. We thank Juliette Meyer from the Bioinformatic hub of Institut Pasteur for advice on statistical analysis. Several bioinformatic analyses were performed on the Core Cluster of the Institut Français de Bioinformatique (IFB) (ANR-11-INBS-0013). We are also grateful to Sandra Legout and the team managing scientific archives at the Institut Pasteur for their kind help in naming the phage collection.

## Data availability

All the data and the code used to generate the analyses presented here is available on the Github repository of the project (https://github.com/mdmparis/coli_phage_interactions_2023). Raw images of the plaque assay experiments are available on the Zenodo repository of the project. The encoded interaction matrix is also available in the Viral Host Range database (https://viralhostrangedb.pasteur.cloud/) [68].

