## Supplementary Materials for "Predicting phage-bacteria interactions at the strain level from genomes"

**Supplementary Tables**

- Supplementary Table 1 : *Picard* Collection
- Supplementary Table 2 : *Guelin* Collection
- Supplementary Table 3 : Interaction matrix
- Supplementary Table 4 : Outer membrane proteins of strains from the *Picard* collection
- Supplementary Table 5 : Defense systems of strains from the *Picard* collection
- Supplementary Table 6 : Test collection
- Supplementary Table 7 : Phage cocktails recommended to the test collection

### Supplementary Texts

#### Supplementary Text 1 – Short biography of Bertrand Picard

Born in 1950, Bertrand Picard is a French medical microbiologist who devoted most of his career in studying *Escherichia coli* natural isolate diversity and the link between strain genetic features and lifestyle.

He obtained his MD degree from the University Paris VII in 1978, followed by a PhD in Biochemistry and Molecular Biology in 1990 at the University Paris V. He completed the molecular taxonomy course at the Institut Pasteur in 1988. He spent most of his hospital career as a medical bacteriologist at the Assistance Publique-Hôpitaux de Paris as Assistant (1980), then Associate (1986) and finally Full Professor (1994), with a 10 year-period in the University hospitals of Brest in Britany.

His research was based on two main intricate objectives. First, he characterized the structural basis of the polymorphism of the carboxylesterase B of *E. coli*. Second, by studying numerous natural isolates from different contexts (commensal, pathogen), in different hosts (humans, non-human mammals, birds) from different countries, he looked for genetic variants associated with a specific lifestyle. His major contribution was to show that strains exhibiting a carboxylesterase $B_2$ electrophoretic type were more frequently found in extraintestinal infections, exhibited numerous virulence associated genes and were highly virulent in a mouse model of sepsis. He is the author of over 100 publications in peer-reviewed journals.

#### Supplementary Text 2 – Short biography of Antonina Guelin

Antonina Guelin (born Chtdedrina) was a French medical doctor and biologist of Russian origin. Born in 1904, in Saint-Petersburg (Russia), she died in 1988. She was first Assistant at the Embryology Laboratory of the Collège de France (1931-1934) where her work focused on the chicken embryo. She then entered the laboratory in the anaerobes department (1934-1941), and the bacteriophage department (1941-1968) headed by Eugène Wollman at the Pasteur Institute. Due to her early works on anaerobic strains, she pioneered the research on phage of *Clostridium perfringens* and *C. histolyticum*. During this period, she notably demonstrated the benefits of monitoring bacteriophages to assess the sanitary quality of fresh water, and the role of bacteriophages in regulation of bacterial population in aquatic environments. For those works, she received an award from the French National Academy of Medicine. She finished her career as a researcher at the Roscoff Biological Station in Britany (France) (1970-1988).

#### Supplementary Text 3 – Link between phage genome size and number of lytic interactions within phage morphotypes

We wondered if the tendency of *Myoviridae* to have a broader host range may be due to their larger genome allowing them to adapt to more diverse host genetic backgrounds at any step of their life cycle (adsorption to host, intracellular replication, etc.). However, when taking the phage morphotype into account, phage genome size did not show a significant association with the number of lytic interactions performed by each phage (Linear model : genome size =2.3x10-4, p=0.17).

#### Supplementary Text 4 – Nestedness and modularity properties of the interaction matrix

A frequently discussed macroscopic property of bipartite networks like phage-bacteria interaction matrices is how the interactions are distributed within the network: from modular (independent groups of phages infect independent groups of bacteria) to nested (host range of more specific phages/bacteria tend to be subsets of the host range of more generalist ones) [15], [94]. Our interaction matrix exhibited strong nestedness (BINMATNEST algorithm, T=11.63, $p<10^{-5}$) and weak modularity (BRIM algorithm, Q=0.23, $p<10^{-3}$) (**Supplementary Figure 2**).

**Supplementary Text 5 - Link between the number of lytic interactions performed by a phage and the number of RBPs encoded in its genome**

We then tested whether the number of lytic interactions performed by a phage host range would be the consequence of the number of RBP it encodes in its genome. Under this hypothesis, we would expect that phages which perform more lytic interactions (e.g. Myoviridae which perform up to 40% of lytic interactions on the *Picard* collection) would have more RBPs than phages which are specific (e.g. Podo- and Siphoviridae which sometimes perform less than 10% of lytic interactions). We ran a systematic analysis on the whole *Guelin* collection of phages and observed that all phages in the collection encode either one or two predicted RBP (See methods) [28]. This estimate was supported by the analysis of the literature in which we found no examples of *Caudoviricetes* with more than two RBP-encoding genes, with the notable exception of phages from the *Ackermannviridae* family harboring four tailspikes in star-like structures [50]. None of these phages were included in the *Guelin* collection.

As a consequence, we did not find any significant correlation between the number of RBP of a phage and the number of lytic interactions it performs (Linear Mixed Model (LMM), $\beta_{Num\ RBP}$= 28.11, 95% credible interval = [-33.09, 83.58], pMCMC = 0.33). This implies that the number of lytic interactions a phage performs is primarily influenced by the tropism of its RBP rather than the number of RBP it encodes.

**Supplementary Text 6 - Multivariate analysis of the bacterial traits associated with lytic interactions in the matrix**

To assess the statistical association between bacterial traits and lytic interactions in the matrix, we also followed a multivariate approach : instead of treating each phage infection separately we fitted a single model on all the phage-bacteria interactions in the matrix. We first fitted a PERMANOVA model to which we provided the presence/absence of both defense system subfamilies and adsorption factors as covariates ("full" model). We compared this full model to (i) a submodel having only adsorption factors as covariates and (ii) a submodel having only defense system subfamilies as covariates. Model comparison was performed using the Akaike Information Criterion (AIC) (See methods). The best possible fit was obtained with the adsorption factors-based model (PERMANOVA, AIC=-435, adjusted $R^2$=0.27) compared with the full model (AIC=-418, adjusted $R^2$=0.28) or the defense systems-based model (AIC=-372, adjusted $R^2$=0.05). This approach also suggests that adsorption factors better explain bacterial host range than bacterial defense mechanisms.

**Supplementary Text 7 – Link between bacterial susceptibility to phages and bacterial genome size or number of antimicrobial genes encoded in the genome**

AMR genes were chosen as a control variable as they undergo evolutionary dynamics similar to the ones of defense systems. Genome size was chosen as it is related with the composition of the pangenome in a given strain [33]. None of these variables were *a priori* expected to be associated with susceptibility to phages but if they were, this would suggest that the observed correlation between the MLC given phage infection and the number of defense systems per genome is more likely to be a "pangenome-related" effect than a true effect of the presence of defense systems (**Supplementary Figure 6**). Only genome size was weakly but significantly associated with the average MLC given infection (Genome size : $\beta_{genome\ size}$ = -4.49x10$^{-7}$, 95% credible interval = [-8.5x10$^{-7}$, -1.0x10$^{-7}$], pMCMC = 0.05 ; AMR genes : $\beta_{Num\ AMR\ genes}$ = -0.03, 95% credible interval = [-0.05, 0.01], pMCMC = 0.11). However, this correlation between the average MLC given infection and the bacterial genome size did not hold once the extreme values in terms of genome size were removed from the dataset. Furthermore, when putting both genome size and the number of defense systems as covariates in a model, only the number of defense systems was significantly associated with the MLC given infection. This indicates that the information carried by the genome size variable was recapitulated by the number of defense systems and that the best fit is obtained with this variable.

### Supplementary Figures

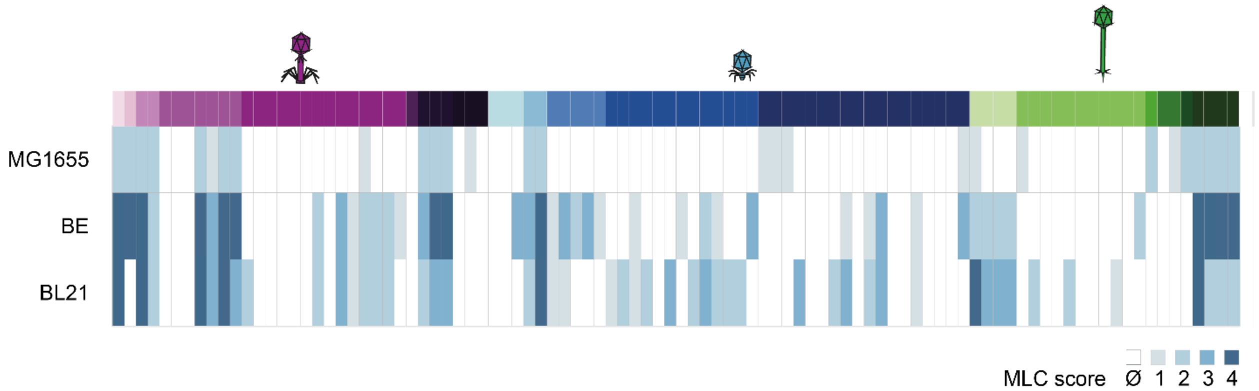


**Supplementary Figure 1. Interactions between the three laboratory-adapted strains MG1655, BE and BL21 with the phages of the *Antonina Guelin* collection.**

Interactions between MG1655 and the collection were performed at single phage dilution (MOI > 1). Consequently, it is only possible to state whether a phage does not perform a lytic interaction (MLC score = Ø), performs lytic interactions with countable lysis plaques (MLC score = 1) or full clearing (MLC score = 2) at MOI = 10. For BE and BL21, the interactions were tested at all phage dilutions so the MLC score ranges from Ø to 4.

A

B

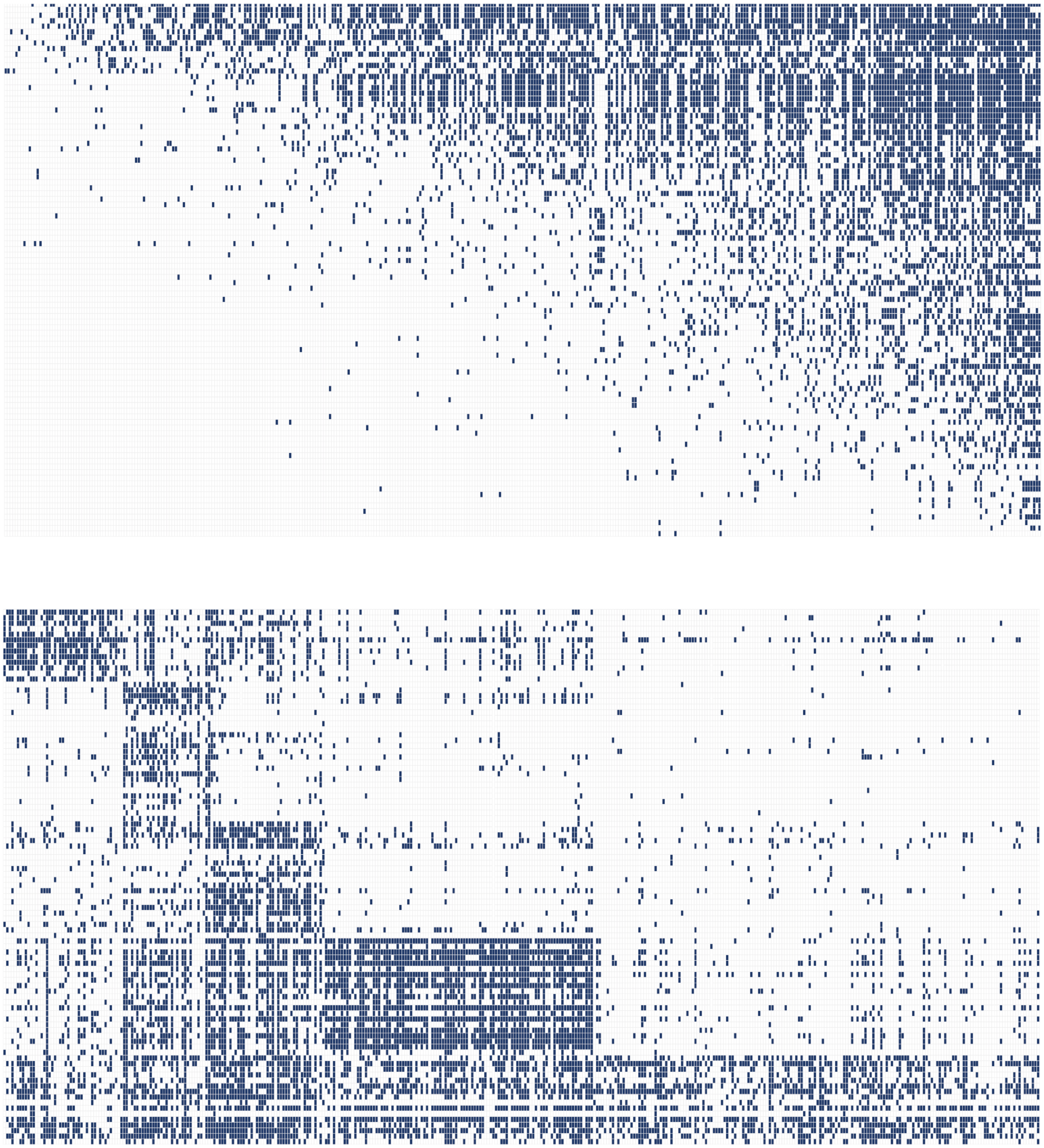

**Supplementary Figure 2. The phage-bacteria interaction matrix is nested and weakly modular.**
**A.** Rows and columns of the interaction matrix are rearranged using the BINMATNEST algorithm [89] in order to exhibit the matrix nestedness. Nestedness indicates that specific phages tend to infect the same bacteria as more generalist ones. The p-value is computed by generating 1,000 random matrices with the same size and average connectance as the original interaction matrix. **B.** Rows and columns of the interaction matrix are rearranged using the BRIM algorithm [90] to exhibit the matrix modularity. As in **A**, the p-value for the BRIM score is computed based on the generation of 1,000 null matrices with similar properties (size and connectance) as the original interaction matrix.

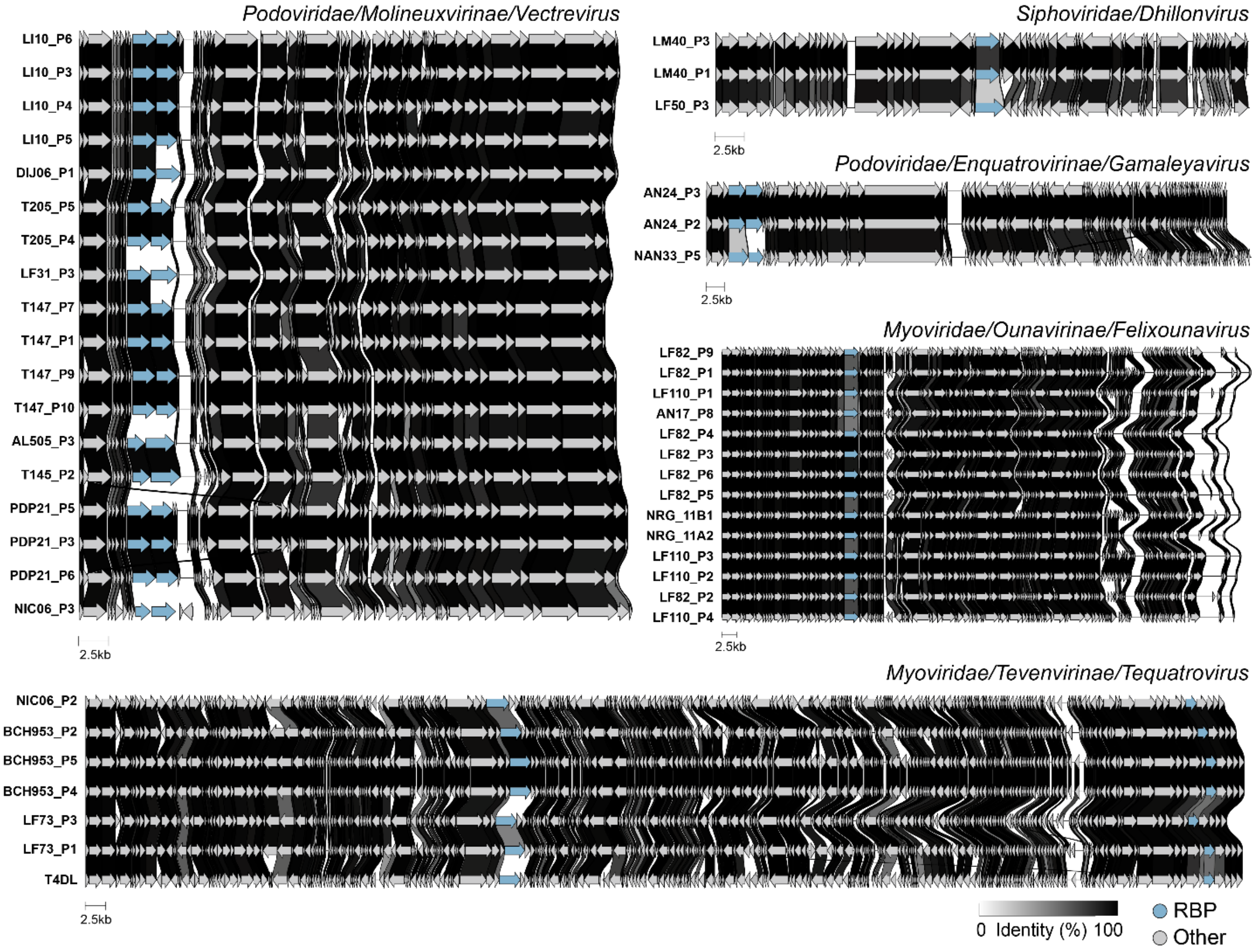


**Supplementary Figure 3. RBP-encoding genes are hotspots of intra-genus genomic variability.** Whole genome pairwise alignment of phage genomes belonging to the same genus, on five example genera (*Vectrevirus*, *Dhillonvirus*, *Gamaleyavirus*, *Felixounavirus* and *Tequatrovirus*) [56]. The stroke between two subsequent open reading frames (ORF) at the same locus indicates the percentage of amino acid identity between them. ORFs encoding for RBP are highlighted in light blue.

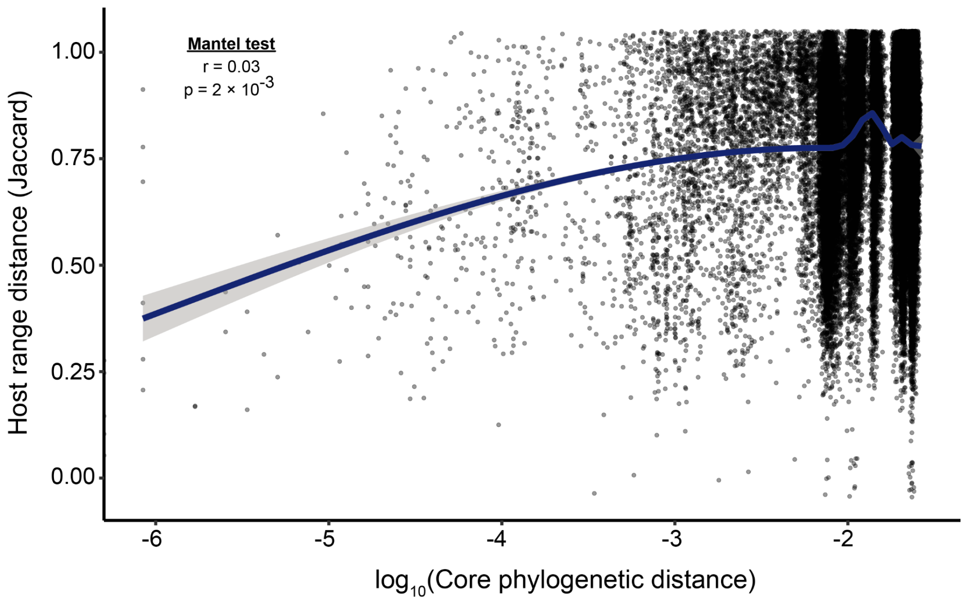


**Supplementary Figure 4. Bacterial core phylogeny weakly correlates with bacterial infection patterns.**

For each possible couple of bacterial isolates, the core phylogenetic distance is computed [83] as well as the distance between their respective host ranges. For the latter, the host range of each isolate is binarized (null MLC score *vs.* positive MLC score) and the Jaccard distance is used. Mantel test is used to assess the correlation between each set of distances [47]. The best fit cubic spline is shown (blue).

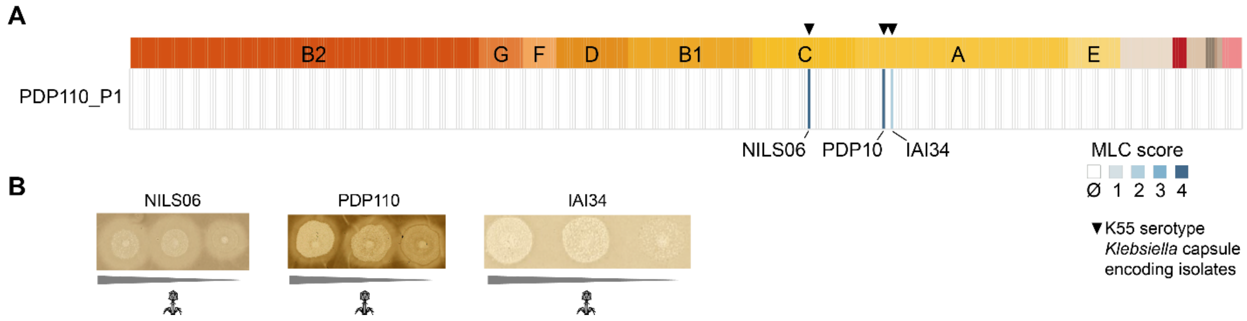


**Supplementary Figure 5. PDP110_P1 is likely to be a *Klebsiella* capsule K55-targeting phage.**
**A. PDP110_P1 only infects three strains in the *Picard* collection.** Interactions between phage PDP110_P1 (*Siphoviridae/Lambdavirus*) and the *Escherichia* isolates belonging to the *Picard* collection. The name of each isolate infected by PDP110_P1 is provided. Isolates which encode a K55-serotype *Klebsiella* capsule are indicated (black triangle). **B. PDP110_P1 tends to generate halos on the strains it infects.** Raw plaque assay results are shown for the three isolates which are infected by PDP110_P1 (NILS06, PDP110 and IAI34). The interactions are tested at three phage MOI (10, 1 and 0.1 from left to right).

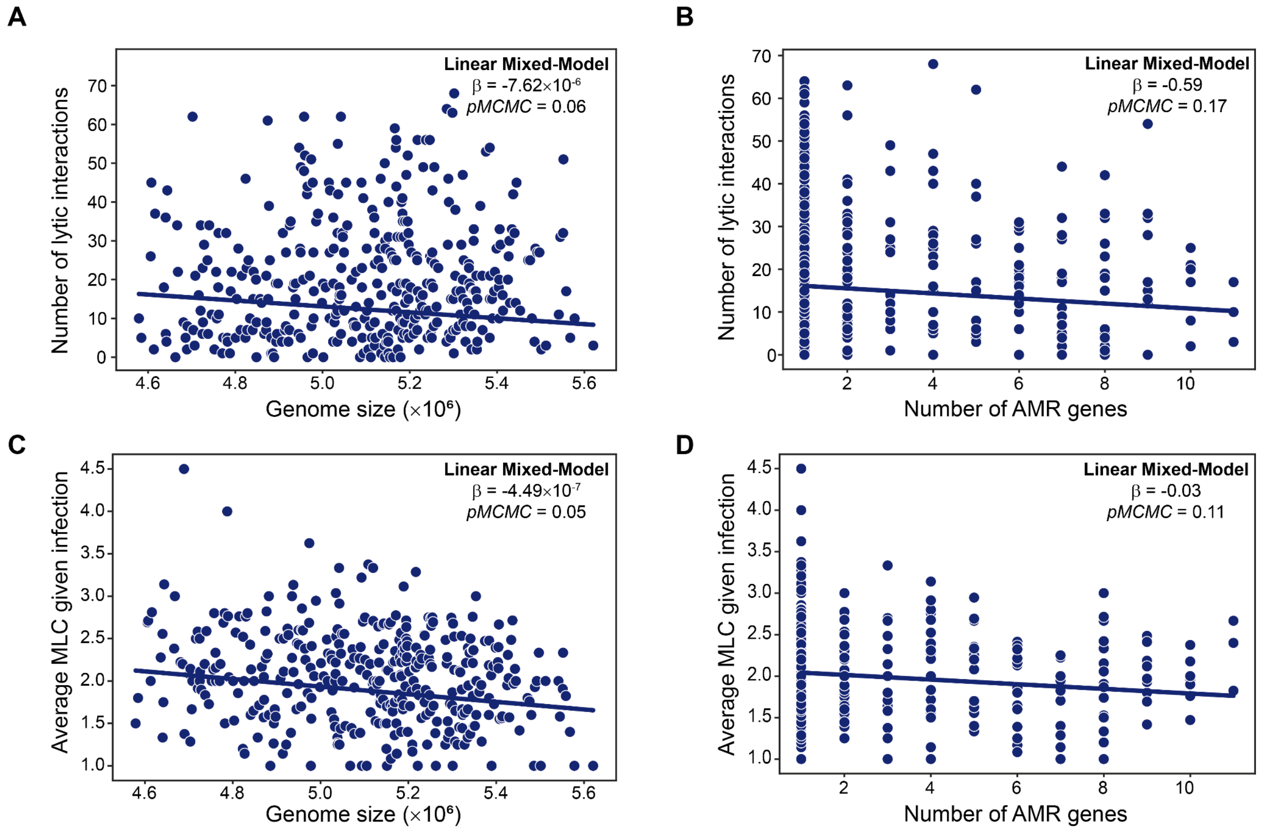


**Supplementary Figure 6. Bacterial Genome size significantly correlates with phage virulence upon infection, but the number of antimicrobial resistance (AMR) genes encoded in the bacterial genome does not.**

A LMM is fitted to the number of lytic interactions undergone by each bacterial strain (**A-B**) or the average MLC upon infection (**C-D**), taking as a covariate either the bacterial genome size (**A-C**) or the number of antimicrobial resistance genes encoded in the bacterial genome (**B-D**). The LMM takes the inverse relatedness matrix computed from the bacterial core phylogenetic distance as a random effect [57]. Significance is assessed based on the pMCMC which gives the probability of the estimated coefficient being null under the estimated posterior distribution.

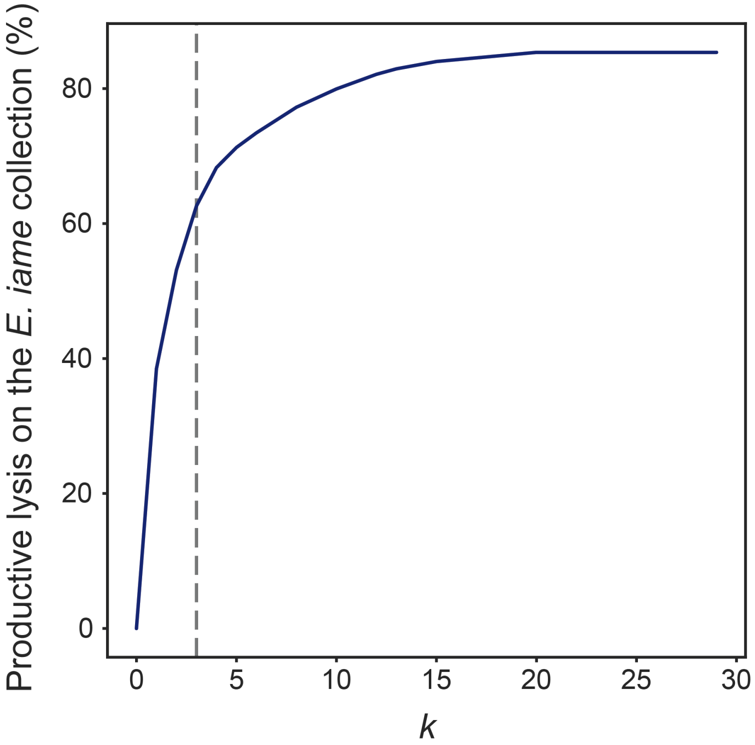


**Supplementary Figure 7. Coverage of the *Picard* collection by the most infecting *k*-uplet of phages of the *Antonina Guelin* collection.** The number of productive lysis events performed by the most covering *k*-uplet of phages is computed for diverse values of *k* from 1 to 25 (blue). The gray dashed line corresponds to the most covering triplet (*k* = 3) of phages, which was chosen as a naive baseline to compete with our phage cocktail recommender algorithm. This triplet performs 63% of productive lysis on the *Picard* collection.

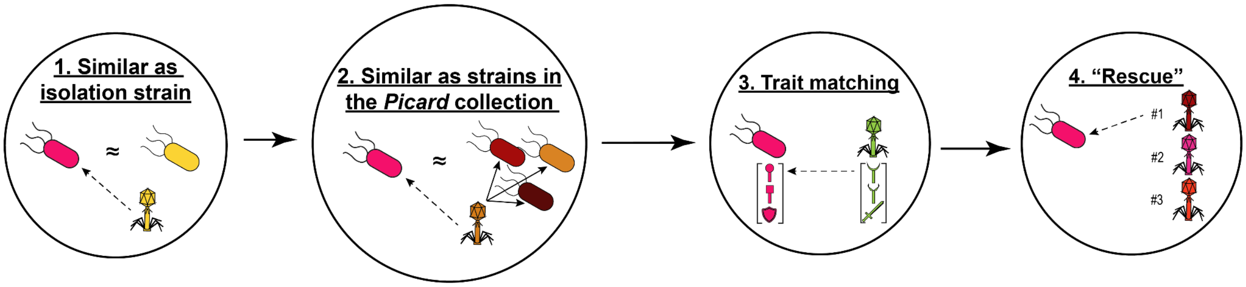


**Supplementary Figure 8. A four-steps algorithm to recommend phage cocktails targeting *Escherichia* strains.** The recommender algorithm for phage cocktails is designed as a four-steps pipeline. Each step aims at determining whether each phage in the *Guelin* collection should be added to the recommended cocktail regarding a specific criterion which is defined by the input features and the algorithm that is used. The algorithm goes sequentially through the four steps and stops whenever $k$=3 phages are recommended. In the recommendation process, the pipeline is forbidden to recommend two phages from the same genus and same isolation strain to ensure phage diversity within the cocktail (See methods).

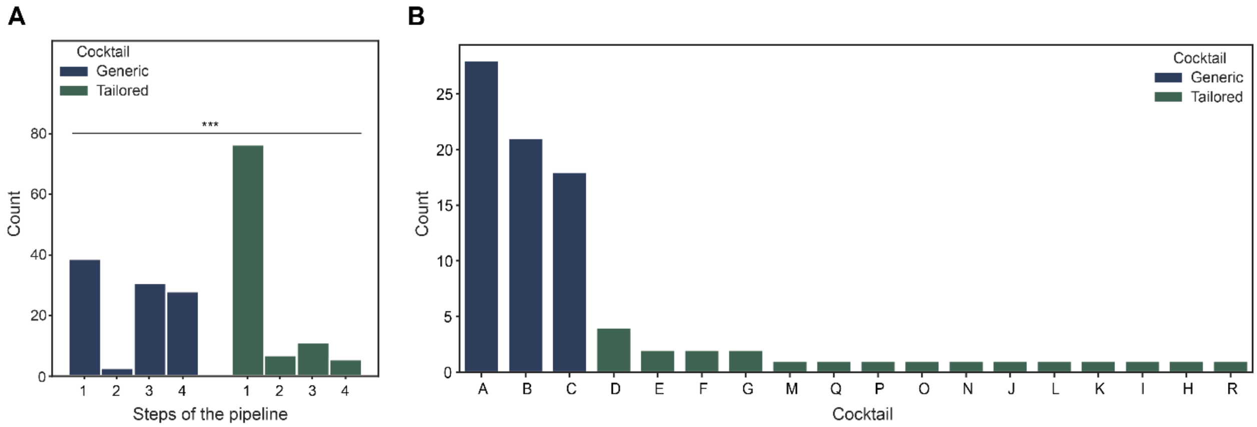


**Supplementary Figure 9. Recommended cocktails can be partitioned into generic and tailored cocktails.**

**A. Distribution of the steps of the pipeline at which each phage is recommended in the pipeline.** We recommend phage cocktails composed of three phages to each of the 100 test strains. The recommender algorithm goes sequentially through four steps. This design choice causes the algorithms to be partitioned into two categories : "Generic" and "Tailored". The distribution of the steps of the pipeline at which phages are recommended depending on the cocktail category is represented. *** : $p < 10^{-3}$ (two-sided Mann-Whitney test). **B. Number of haplotypes to which each cocktail is recommended.** The haplotype of a bacterial strain is defined by the combination of its Sequence Type (ST), O-antigen serotype and H-antigen serotype. The number of haplotypes is a measure of the genetic diversity of the bacteria which are recommended a specific cocktail. Cocktails are partitioned between "Generic" and "Tailored" cocktails.
